# Programmed DNA breaks drive chromatin reconfiguration and facilitate transcriptional potentiation at neuronal early response genes

**DOI:** 10.1101/2025.05.06.652469

**Authors:** Lance Heady, Richard Rueda, Amir Segev, Krystal G. Morton, Ram Madabhushi

## Abstract

Neuronal activity causes topoisomerase IIβ (TOP2B) to form DNA double strand breaks (DSBs) within the promoters of key early response genes (ERGs), such as *Fos* and *Npas4*. TOP2B-mediated DSBs facilitate rapid ERG transcription, yet how this occurs remains unclear. Here, using chromosome conformation capture methods (3C and 4C-seq), we report that DSB formation within the promoters of *Fos* and *Npas4* is sufficient to emulate contact profiles observed at these regions following neuronal stimulation, including their elevated interactions with cognate enhancers. Furthermore, despite their purported risk, repeated DSB cycles within ERG promoters progressively potentiated ERG induction in both mouse cortical neurons and HEK293T cells, evoking the effects of transcriptional memory. Potentiated ERG inducibility following recurrent DSBs persisted through intervening cell cycles, occurred even when DNA repair was likely mutagenic, and was associated with a substantial loss of *cis* chromosome interactions and an increase in *trans* interactions with ERG promoters. Together, these results reveal how single and recurrent TOP2B-mediated DSBs could affect stimulus-dependent transcription patterns by affecting chromatin dynamics at ERG promoters.

## INTRODUCTION

Stimulus-dependent transcription programs in neurons allow animals to alter neural connectivity and develop adaptive responses to new experiences, such as forming long-term memories^1–5^. Following neuronal stimulation, genes aptly named early response genes (ERGs) are rapidly transcribed within the first few minutes and are enriched in transcription factors, such as *Fos, Npas4, FosB,* and *Egr1*^5–7^. ERG products then prime the transcription of late response genes (LRGs) that display cell type-specific expression patterns and functions, including mediating experience-driven synaptic changes^5,6^. Disruptions in neuronal activity-induced transcription manifests in various intellectual disability and autism spectrum disorders^8^. Understanding how activity-induced transcription is controlled is therefore important.

Whereas the hallmark of ERGs is their rapid activity-dependent transcription, how this occurs is unresolved^7^. ERG promoters are pre-bound by activity-responsive transcription factors and by RNA polymerase II (RNAPII), which initiates transcription but is paused at promoter-proximal regions in unstimulated neurons^9^. Activity-induced synapse-to-nucleus signaling then sparks the escape of paused RNAPII to drive ERG transcription. Although how RNAPII pause-release is achieved remains unclear, previous reports indicate that this step requires interactions between ERG promoters and cognate enhancers^9–13^. For instance, sequences within the promoter of the ERG, *Arc*, activate transcription from *Arc* enhancers and these enhancer RNAs (eRNAs), in turn, are thought to mediate RNAPII pause-release from the *Arc* promoter^9,10^. Distinct combinations of enhancers interact with ERG promoters following neuronal activity and these contact patterns confer robust and stimulus-specific ERG transcription^12,13^. Overall, these results suggest that mediators of enhancer-promoter interactions could comprise the key trigger for rapid ERG transcription.

Yet how ERG promoters and often distantly located enhancers contact each other as a function of neuronal activity is ill-defined. For instance, recent reports indicate that structural maintenance of chromosomes (SMC) complexes, such as cohesin, reel and extrude chromatin to generate loops^14–20^. Whereas loop extrusion can bring distal chromatin regions into proximity, cohesin loss affected neither activity-induced enhancer-promoter contacts nor the induction of neuronal ERGs^21^. These results imply the existence of other mechanisms that orchestrate activity-dependent communication between gene regulatory elements in neurons.

We previously reported that various paradigms of neuronal activity, including the exposure of mice to learning behaviors, induce the topoisomerase, TOP2B, to generate DNA DSBs at specific genomic sites^22^. These loci are enriched for the promoters of neuronal ERGs and DSB formation in this manner facilitates rapid ERG transcription^22,23^. DSBs have been linked to stimulus-dependent transcription in the brain as well as in diverse cell types following various stimuli^22–30^. Despite this, how activity-induced DSBs facilitate the transcription of associated genes, including ERGs, is poorly understood. TOP2B was shown to promote RNAPII pause-release from ERG promoters, which is rate-limiting for ERG transcription^24^. Furthermore, ERG promoters are bound by the architectural protein, CTCF, which has known enhancer-blocking activity and TOP2B-mediated DSBs form in the vicinity of CTCF binding sites in neurons^22,31,32^. Based on these results, we hypothesized that DSBs could induce rapid transcription by overriding enhancer-blocking chromatin topologies at ERG promoters.

To test this idea, here we utilized circular chromosome conformation capture sequencing (4C-seq)^33,34^ to assess genome-wide interactions at the promoters of the ERGs, *Fos* and *Npas4*, as a function of neuronal stimulation and TOP2B-mediated DSB formation. We find that neuronal stimulation increases contacts of ERG promoters not just with putative enhancers but with regions broadly across the *cis* chromosome. Artificially generating DSBs within ERG promoters was sufficient to mimic contact patterns observed following neuronal stimulation. These results suggest that DSBs drive transcription by increasing chromosome dynamics at ERG promoters. Activity-induced DSBs have often been considered a risky event because inaccurate DSB repair could lead to mutation accrual and compromise ERG transcription upon subsequent rounds of neuronal stimulation^25,35,36^. Unexpectedly, recurrent DSB formation within ERG promoters progressively potentiated ERG transcriptional induction both in neurons and in HEK293T cells in a manner indicative of transcriptional memory. Potentiated ERG inducibility in both neurons and HEK293T cells correlated with the pruning of *cis* chromosome interactions and an increase in *trans* interactions with ERG promoters. Overall, these observations reveal how TOP2B-mediated DSBs impact stimulus-driven transcription programs by rewiring chromosome interactions at ERGs.

## RESULTS

### TOP2B-mediated DSBs stimulate enhancer-promoter interactions at the neuronal *Fos* locus

Chromosome conformation capture experiments (3C and 5C) indicate that neuronal stimulation increases contacts between ERG promoters and specific enhancers^12,13^. To understand how TOP2B-mediated DSBs affect enhancer-promoter communication at ERGs, we treated cultured primary mouse cortical neurons (days *in vitro* 13 (DIV13)) with the TOP2 poison, etoposide (ETP; 10 μM for 50 min), which induces TOP2B to generate DSBs within the promoters of ERGs, including *Fos*, in the absence of neuronal stimulation^22,23^. Separately, cultured neurons were stimulated by brief incubation with N-methyl-D-aspartate (NMDA; 50 μM for 10 min followed by washout for 20 min), which results in activity-induced DSB formation by TOP2B^22,23^. Quantitative real-time PCR experiments verified that the expression of *Fos* and other ERGs was markedly upregulated in NMDA-treated neurons and modestly upregulated in ETP-treated neurons (Figures S1A and S1B)^22,23^. Pairwise interactions of the *Fos* promoter with its previously annotated enhancers (E1 and E2) were then assessed using 3C-qPCR in untreated, NMDA-treated, and ETP-treated neurons^13^. As reported for neurons stimulated with either KCl or bicuculline, interactions between the *Fos* promoter and enhancer, E2, were elevated (1.67-fold) while interactions with E1 were unaffected, in NMDA-treated neurons relative to unstimulated controls (Figure S1C)^12,13^. Notably, ETP treatment also caused a two-fold increase in E2-promoter contacts and a trending increase in E1-promoter contacts at *Fos* (Figure S1C). These results suggest that TOP2B-mediated DSB formation is sufficient to trigger enhancer-promoter contacts at ERGs that are observed in response to neuronal activity.

### TOP2B-mediated DSBs increase chromosome dynamics at ERG promoters

Whereas 3C only measures pairwise interactions between known genomic elements, 4C-seq detects genome-wide contacts with a region of interest^33,34^. To explore interactions at sites of activity-induced DSBs in an unbiased manner, we performed 4C-seq using the promoter segments of either *Fos* or *Npas4* as bait and generated normalized contact profiles at these ERG promoters in untreated, NMDA-treated, and ETP-treated neurons. To annotate interacting regions, available chromatin immunoprecipitation sequencing (ChIP-seq) data of key histone marks and ChromHMM were used to first classify mouse neuronal chromatin into 15 states, and then re-grouped into 4 functional classes (promoters, enhancers, gene bodies, and heterochromatin) (Figure S1D)^37–39^. Additionally, available ATAC-seq and transcriptomic data were used to label putative regulatory elements in stimulated and ETP-treated neurons (Figures S1E – S1H)^9,38,40^. 4C-seq contacts were then analyzed at various scales of distance from each bait region (viewpoint), as described below.

As a first step, normalized 4C-seq signals were compared across 300 kb windows surrounding the *Fos* and *Npas4* viewpoints^34^. Chromosome interactions of the two ERG viewpoints were altered with regions across the 300 kb window in neurons treated with either NMDA or ETP (Figures S1E – S1H). For instance, contacts between the *Fos* promoter and several enhancers were increased in NMDA- and ETP-treated neurons relative to controls, while contacts with other enhancers were reduced (Figures S1E and S1G). Similar patterns were observed with contacts at the *Npas4* promoter (Figures S1F and S1H). Interaction changes at both viewpoints were also not limited to enhancers but were detected at various promoter, gene body, heterochromatin, and accessibility states (Figures S1E – S1H). For instance, ETP caused increased contacts between ERG promoters and regions of low chromatin accessibility (Figures S1G and S1H). Overall, NMDA and ETP seemed to increase chromosome dynamics at ERG promoters without a clear preference for contacts based on histone modifications, chromatin accessibility or transcriptional activity.

Next, to assess long-range contacts with ERG promoters, we utilized 4Cker, a Hidden Markov Model-based pipeline that uses adaptive windows to account for the decay in 4C coverage with increasing distance from the viewpoint^41^. 4Cker employs distinct programs to call regions of significant interaction that reside either within 2Mb from the viewpoint (nearbait) or on the same chromosome as the viewpoint (*cis*)^41^. 4Cker analysis revealed that the nearbait interactome at the *Fos* viewpoint, defined as the total coverage of all significant called interacting regions within 2Mb surrounding the viewpoint, increased in size by 100.9 % in NMDA-treated neurons and by 45.1 % in ETP-treated neurons relative to controls (Figure 1A). At the *Npas4* viewpoint, the nearbait interactome increased by 271.4 % in NMDA-treated neurons and by 77.1 % in ETP-treated neurons (Figure 1B). Similarly, NMDA and ETP caused an increase in the *cis* chromosome interactome compared to controls at both viewpoints (Figures 1C and 1D). To further assess the nature of interactions with ERG promoters, ChromHMM was used to segment chromosomes 12 and 19 according to the 4 functional classes of chromatin states, and the number of regions with mapped 4C-seq reads was quantified for each replicate per condition. The number of regions contacted by both ERG viewpoints increased across all chromatin states in NMDA-treated and ETP-treated neurons compared to controls (Figures 1E). These results suggest that TOP2B-mediated DSB formation correlates with increased chromosome-wide contacts at ERG promoters in both stimulated and unstimulated neurons.

**Figure 1:**
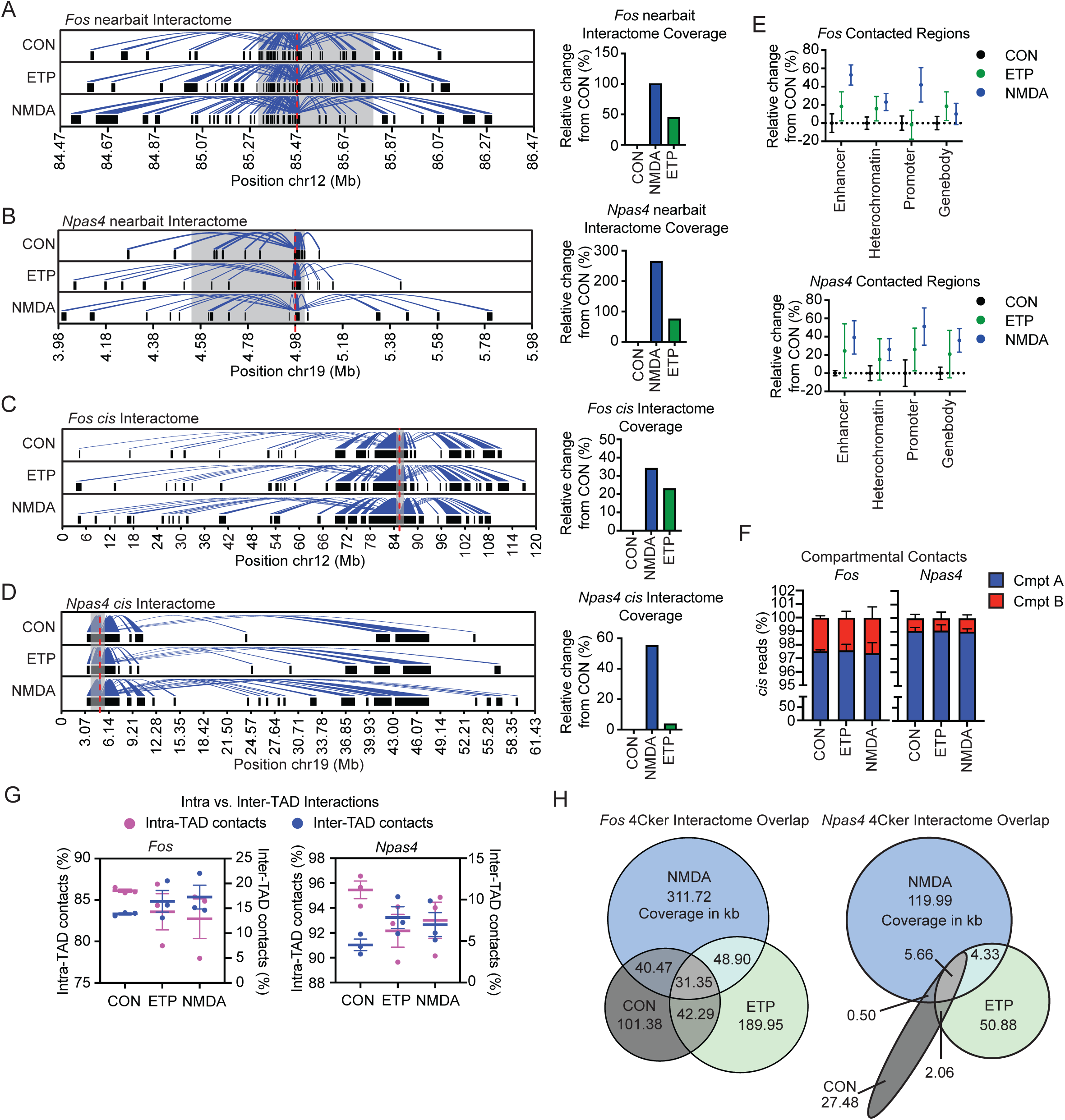
TOP2B-mediated DSBs increase interactions at ERG promoters. **(A-B)** (left) 4Cker was utilized to call significantly interacting regions in CON-, ETP-, and NMDA-treated neurons across three 4C-seq replicates in the nearbait setting (regions within 2MB of the viewpoint). The called interactions (the interactome) are shown via the black boxes which represent regions in the nearbait view that have significant interaction with the viewpoint (shown as the red dotted line). The interaction or looping is shown via arc plots from the viewpoint to each called region (blue arc lines). Previously published Hi-C data was utilized to call TAD boundaries, and the region between the closest TAD boundary upstream and downstream of the viewpoint is shaded in light grey. The called nearbait interactome of untreated (CON), NMDA-, and ETP-treated neurons are shown at both the *Fos* (A) and *Npas4* (B) viewpoints. (right) The coverage of the nearbait interactome for each treatment was calculated in bp and the relative change in size from untreated neurons (CON) is shown for each viewpoint. **(C-D)** 4Cker was utilized to call significantly interacting regions in CON-, ETP-, and NMDA-treated neurons across three 4C-seq replicates in the *cis* setting (all regions on the *cis* chromosome) The called interactions (the interactome) are shown via the black boxes which represent regions in the on the *cis* chromosome that have significant interaction with the viewpoint (shown as the red dotted line). Interactions, coverage plots, and TAD boundaries are as in (A-B). **(E)** Chromosomes 12 and 19 were split into non-overlapping, functionally distinct regions via ChromHMM segmentation that produced 15 chromatin states. The 15 states were clustered into four broader classifications: enhancer, heterochromatin, promoter, and gene body states. The number of regions with mapped 4C reads was assessed for CON-, ETP-, and NMDA-treated neurons in each replicate. The average relative change in the number of contacted regions from the average CON level is shown for both the *Fos* (top) and *Npas4* (bottom) viewpoints. **(F)** Previously published Hi-C data from mouse cortical neurons was used to define chromatin compartments. The amount of mapped cis reads that occur in compartment A and B were assessed for each replicate in CON-, ETP-, and NMDA-treated neurons and the average percentage of each are shown via stacked bar graphs (blue – Cmpt A, red – Cmpt B) across treatments for both Fos (left) and Npas4 (right) viewpoints. **(G)** Previously published Hi-C data from mouse cortical neurons was used to identify TAD boundaries at 5kb resolution. The closest boundary elements upstream and downstream of the viewpoint were utilized to define the TAD that each viewpoint is in. Mapped *cis* reads were then split into either intra-TAD interactions (those within the two boundaries elements) or inter-TAD interactions (outside of the defined boundary elements). The percent of intra- and inter-TAD reads out of the total cis read amount were assessed for each replicate in CON-, ETP-, and NMDA-treated neurons. The pink symbols represent the intra-TAD values which are graphed on the left y-axis while the blue symbols represent inter-TAD values which are graphed on the right y-axis. This is shown for both the *Fos* (left) and *Npas4* (right) viewpoints **(H)** The nearbait interactome of CON-, ETP-, and NMDA-treated neurons were intersected in a pairwise manner to determine the amount of shared and unique regions of the called interactome of both the *Fos* (left) and *Npas4* (right) promoters. The coverage was calculated for the overlap and the values, in kb, are shown.

Eukaryotic chromosomes are hierarchically folded into topologically associated domains (TADs), which constrain chromatin interactions through looping^42,43^. Additionally, transcriptionally active and inactive regions partition into chromosome compartments A and B, respectively, and compartments preferentially engage in homotypic interactions (A-A and B-B contacts are preferred over A-B)^43,44^. To understand how chromosome organization at these levels affect interactions with ERG promoters, Hi-C data from mouse cortical neurons was used to identify TADs and chromosome compartments^45,46^. The distribution of mapped *cis* reads within each compartment and as a function of TAD organization was then assessed. The ratio of contacts in A and B compartments were unchanged across conditions and a majority of interactions with ERG viewpoints occurred with regions in compartments A (Figure 1F). However, both NMDA- and ETP-treated neurons showed increased proportions of inter-TAD contacts with ERG promoters compared to controls (Figure 1G). These results indicate that DSB formation at ERGs helps override insulation conferred by TAD organization.

Finally, overlaying significant interacting regions across conditions revealed that despite a net increase in contacts, treatments with NMDA and ETP also cause substantial decommissioning of interactions detected in untreated controls. For instance, only 33.4 % and 34.2 % of nearbait interactions at the *Fos* viewpoint in untreated neurons were retained in NMDA- and ETP-treated neurons, respectively (Figure 1H). Similar results were observed at the *Npas4* viewpoint (Figure 1H). These results suggest that activity-induced ERG transcription correlates with the rapid remodeling of chromosome contacts with ERG promoters and that TOP2B-mediated DSBs are sufficient to trigger these changes.

### Prior TOP2B-mediated DSB formation potentiates activity-induced ERG transcription

Neurons undergo recurrent cycles of activity-dependent transcription during an animal’s lifetime. While TOP2B-mediated DSBs facilitate the transcription of neuronal ERGs following a single stimulus, their impact in subsequent rounds of ERG transcription remains uncharacterized. To address this issue, cultured cortical neurons were incubated with ETP (10 μM, 60 min) followed by washout and recovery for 23 hours. Thereafter, neurons were treated either with ETP again or with NMDA (Figure S2A). As an additional strategy for TOP2B-mediated DSB formation, we attempted spaced treatments with either KCl or NMDA. However, unlike ETP, both KCl and NMDA proved to be toxic for long-term assessments and were hence not used (Figures S2B – S2D; data not shown). Treatment of neurons with ETP increased total levels of the DSB marker, γH2AX, after 60 min, indicating the formation of TOP2B-mediated DSBs (Figures S2E and S2F). ChIP-qPCR experiments verified that consistent with the formation of DSBs within ERG promoters, ETP treatment caused ©H2AX levels to accumulate within the gene bodies of *Fos* and *Npas4* (Figure S2G)^22^. ©H2AX signals returned to baseline levels after the 23 h recovery period, indicating successful DSB repair (Figures S2E – S2G). Treating neurons with either ETP or stimulating them with NMDA following this recovery period again increased ©H2AX levels (Figures S2E, S2F, S2H and S2I). These results suggest that multiple rounds of TOP2B-mediated DSBs are triggered at ERG promoters in response to our treatments. Based on this, we assessed how a second round of DSB formation affects ERG induction. Surprisingly, both NMDA and ETP caused higher ERG induction in neurons that were previously treated with ETP than in neurons treated singly with ETP and NMDA (Figures 2A and 2B). Thus, in addition to facilitating ERG transcription immediately following neuronal activity, TOP2B-mediated DSBs also potentiated the inducibility of ERGs in the ensuing round of neuronal stimulation.

**Figure 2:**
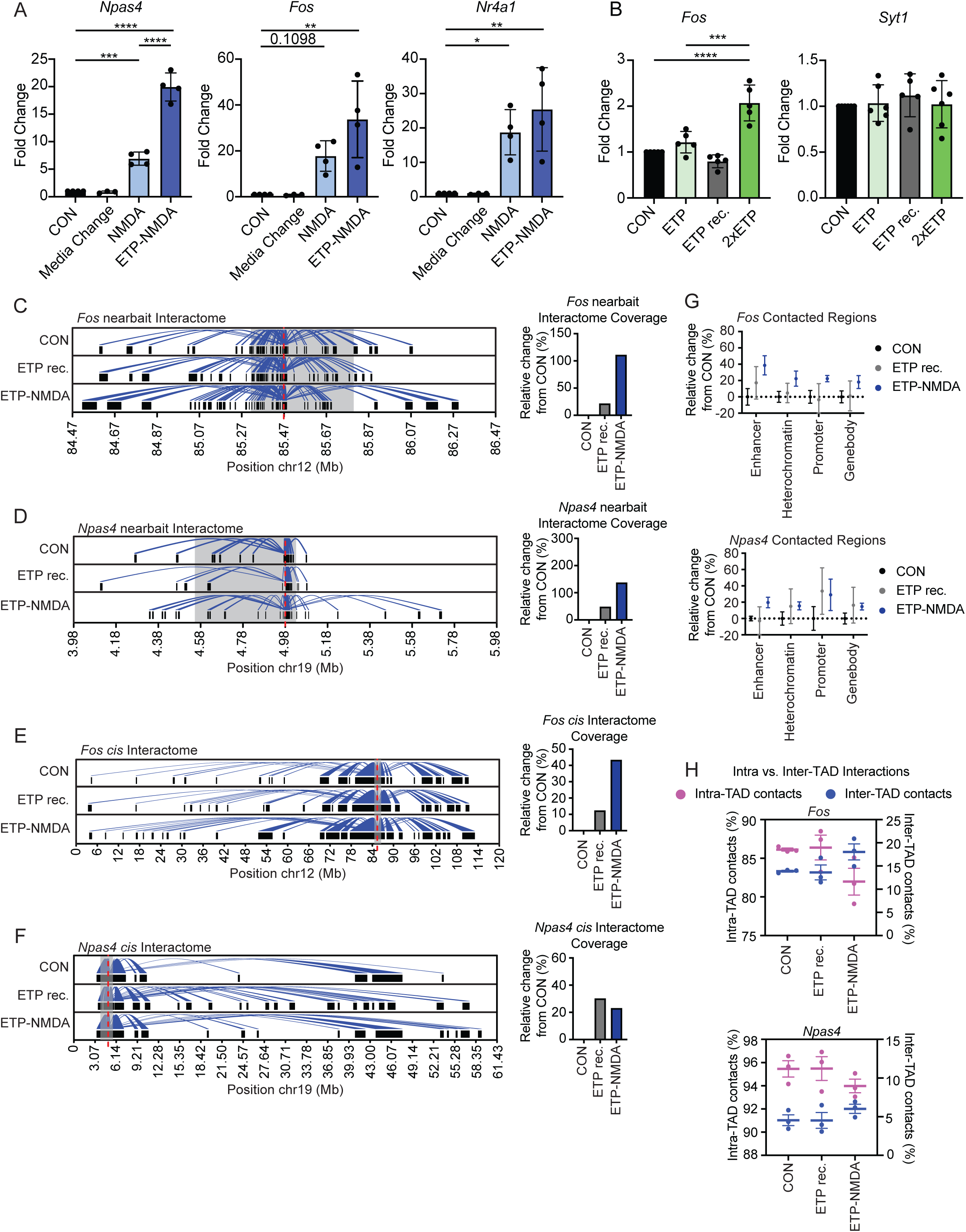
TOP2B-mediated DSBs elevate ERG transcription following an ensuing stimulus. **(A)** Neurons were treated with either NMDA alone or ETP-NMDA and transcript levels were compared to untreated CON-neurons and neurons only subjected to a media change that occurs following repeated stimulations. Total RNA was isolated and gene expression was accessed via RT-qPCR. **(B)** Neurons were treated with either a single round of ETP, and collected immediately, after a 23 hr recovery period, or following a second round of ETP treatment. Gene expression was then accessed via RT-qPCR. **(C-D)** 4Cker, in the nearbait setting (within 2MB of the viewpoint), was utilized to call significantly interacting regions across CON-(same control sample as shown in Figure 1), ETP rec.-, and ETP-NMDA treated neurons in three 4C-seq replicates. Interactions, coverage plots, and TAD boundaries are as described in Figures 1A and 1B. **(E-F)** 4Cker, in the *cis* setting (all regions on the *cis* chromosome), was utilized to call significantly interacting regions in CON-, ETP rec.-, and ETP-NMDA-treated neurons across three 4C-seq replicates. Interactions, coverage plots, and TAD boundaries are as described in Figure 1A and 1B. **(G)** Chromosomes 12 and 19 were segmented as described in Figure 1E. The number of regions with mapped 4C reads was assessed for CON-, ETP rec.-, and ETP-NMDA-treated neurons in each replicate. The average relative change in the number of contacted regions from the average CON (same control sample as shown in Figure 1) level is shown for both the *Fos* (top) and *Npas4* (bottom) viewpoints. **(H)** The percent of intra- and inter-TAD reads (defined as described in Figure 1G) out of the total *cis* read amount were assessed for CON- (same control sample as shown in Figure 1), ETP rec.-, and ETP-NMDA-treated neurons across three 4C-seq replicates. The pink symbols represent the intra-TAD values which are graphed on the left y-axis while the blue symbols represent inter-TAD values which are graphed on the right y-axis. This is shown for both the *Fos* (top) and *Npas4* (bottom) viewpoints.

### Successive rounds of TOP2B-induced DSB formation alter chromosome contact patterns at neuronal ERGs

To understand how chromatin interactions at ERG promoters relate to their potentiated induction following two rounds of DSB formation, we performed 4C-seq in ETP-treated neurons after the 23-hour recovery period (ETP rec.) and after their stimulation with NMDA (ETP-NMDA) (Figure S2A). Normalized contact profiles were again generated at *Fos* and *Npas4* viewpoints for these conditions and compared to those from untreated neurons and neurons treated singly with either ETP or NMDA.

Analysis of 4C signals indicated that several local contacts that formed rapidly upon ETP treatment were maintained even after the cessation of ERG transcription and the repair of TOP2B-mediated DSBs (Figures S3A and S3E). These results suggest that chromatin interactions with ERGs are distinct in ETP rec. neurons than CON neurons at the time of their stimulation of with NMDA. Based on these observations, local 4C signals were compared in ETP-NMDA-treated and NMDA-treated neurons. Unexpectedly, several regions, including enhancers that displayed enriched contacts with ERG viewpoints in NMDA-treated neurons, showed reduced contacts in ETP-NMDA-treated neurons (Figures S3B-S3D, and S3F-S3H). To further understand how chromosome interactions evolve in response to two rounds of DSB formation, 4Cker was again employed to assess genome-wide significant contacts in ETP rec. and ETP-NMDA-treated neurons. Analysis of nearbait interactions with *Fos* and *Npas4* viewpoints revealed an increase in significant contacts in ETP-NMDA-treated neurons compared to ETP rec. neurons (Figures 2C and 2D). For instance, total interactome size at the *Fos* viewpoint increased by 73.9 %, while the interactome at the *Npas4* viewpoint increased by 60.4 %, in ETP-NMDA-treated neurons compared to ETP rec. neurons (Figures 2C and 2D). However, these increases in interactome size were lower than those observed in NMDA-treated neurons compared to untreated controls (100.9% increase at *Fos* and a 271.4 % increase at *Npas4*; Figures 1A and 1B). Similar trends were also observed in relative *cis* interactome size changes at both viewpoints (Figures 2E and 2F).

Additionally, the number of regions that contacted the *Fos* viewpoint were increased in ETP-NMDA-treated neurons compared to ETP rec. neurons and distributed across all 4 classes of chromatin states (Figures 2G). However, this increase was lower than that observed in NMDA-treated neurons compared to untreated controls (Figure 1E). This difference was even more prominent at the *Npas4* viewpoint, where only enhancers showed an increase in contacts with the viewpoint in ETP-NMDA-treated neurons compared to ETP rec. neurons, while other classes either showed no difference or even a slight reduction (Figure 2G). In fact, a reduced number of unique regions contacted both viewpoints in ETP-NMDA-treated neurons when compared to NMDA-treated neurons (1148 vs 1161 for *Fos* and 602 vs 674 for *Npas4*, respectively). Analysis of intra-TAD and inter-TAD interactions at these stages revealed that completion of DNA repair in ETP rec correlated with the reduction of inter-TAD interactions to levels seen in untreated controls (Figure 2H). Inter-TAD interactions were again elevated at both *Fos* and *Npas4* viewpoints following subsequent neuronal stimulation in ETP-NMDA neurons (Figure 2H). These results suggest that neuronal stimulation following a prior round of DSB formation potentiates ERG transcription and is associated with a reduced propensity for *cis* chromatin interactions with ERG promoters.

### *Cis* chromosome contacts with ERG promoters are decreased and contacts in *trans* are increased, following recurrent TOP2B-mediated DSB formation

Transcription and chromosome interaction patterns observed at ERGs following two rounds of DSB formation prompted us to further investigate how these patterns evolve with recurrent DSB cycles. To this end, we performed once-daily ETP treatments starting with DIV9 cultured mouse cortical neurons for 4 consecutive days. Following the 23 hour recovery period (4xETP rec.), neurons were treated either with ETP again (5xETP) or with NMDA (4xETP-NMDA) (Figure S4A). Remarkably, qRT-PCR analysis indicated a progressive potentiation of ERG transcription with recurrent TOP2B-mediated DSB formation (Figures 3A and 3B). These results suggest that DSB formation at ERG promoters is sufficient to elicit transcriptional memory for ERG induction. To determine how chromatin configuration at ERG promoters relates to enhanced ERG induction, we performed 4C-seq in 4xETP rec., 5xETP, and 4xETP-NMDA neurons again using the *Fos* and *Npas4* promoter viewpoints as bait. Comparison of 4C-seq signals revealed reduced contacts in 4xETP rec. neurons compared to untreated controls across the 300 kb window surrounding both viewpoints (Figures S4B and S4H). Thus, while a single round of TOP2B-mediated DSBs caused selective new interactions to be maintained for at least 24 h, recurrent DSB cycles caused the pruning of such interactions. Analysis of 4C-seq signals in 5xETP- and 4xETP-NMDA-treated neurons relative to this new baseline of contacts in 4xETP rec. revealed relatively modest changes in interactions across the 300 kb window with both viewpoints (Figures S4C-S4G and S4I-S4M). For instance, contacts with enhancer, E1, were elevated in 4xETP-NMDA-treated neurons while contacts with enhancer, E2, which showed elevated contacts in NMDA- and ETP-treated neurons, were reduced at the *Fos* viewpoint (Figure S4D, S4F, and S4G). At the *Npas4* viewpoint, contacts with an internal enhancer-like region were gained in 4xETP-NMDA compared to 4xETP rec. (Figures S4J, S4L, and SS4M). Despite these changes, local chromosome contacts with both viewpoints were substantially attenuated in 4x ETP rec., 5xETP, and 4xETP-NMDA neurons compared to untreated controls (Figures S4B-S4G and S4H-S4M)).

**Figure 3:**
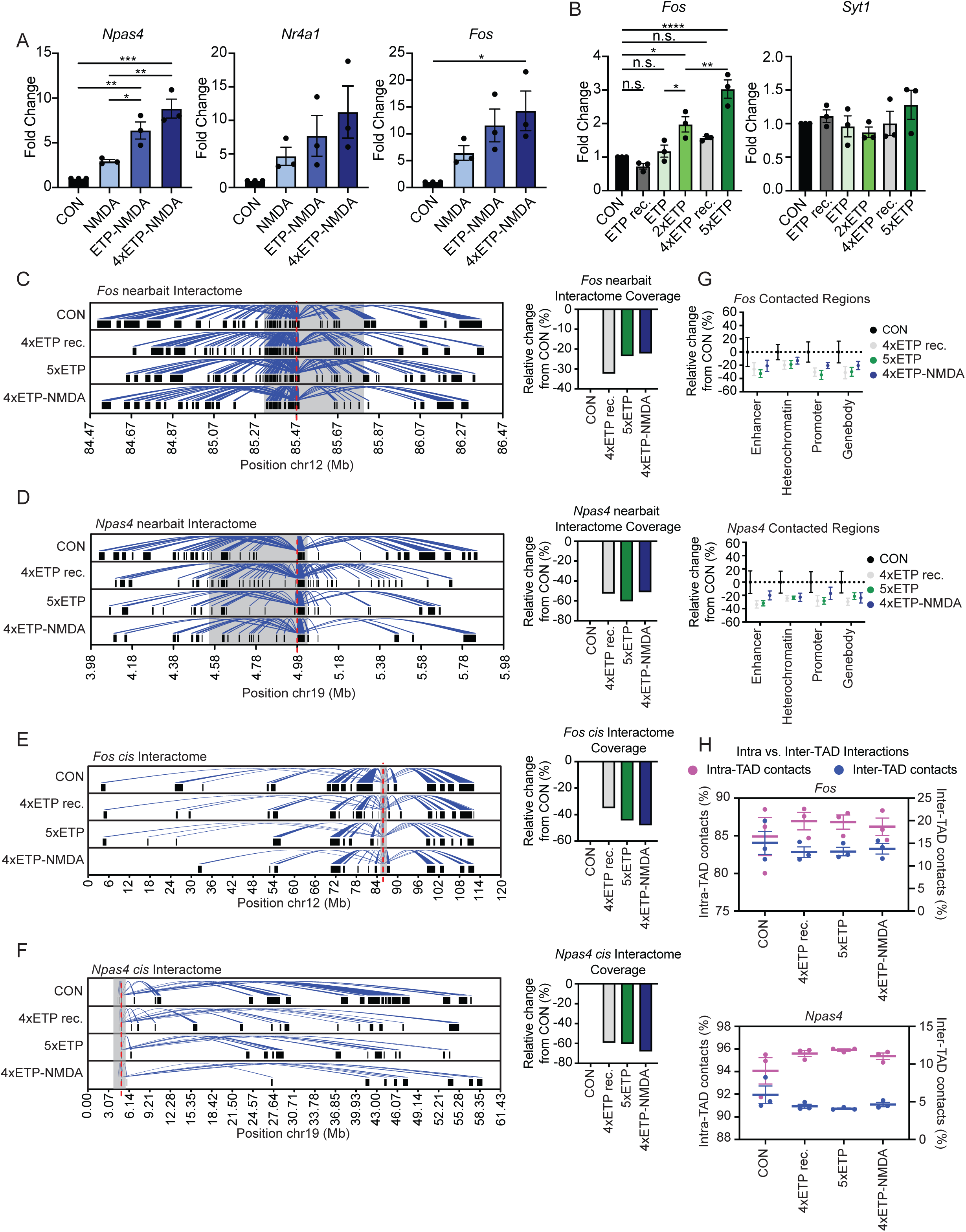
Multiple rounds of TOP2B-mediated break formation cause progressive potentiation of the ERGs and is associated with decreased *cis* interactions. **(A)** Neurons were treated with either NMDA, ETP-NMDA, or 4xETP-NMDA and transcript levels were compared to untreated, CON-neurons. Total RNA was isolated, and gene expression was accessed via RT-qPCR. **(B)** Neurons were treated with either a single round of ETP (collected immediately after stimulation and after a 23hr recovery period), two rounds of ETP-2xETP, four rounds of ETP-4xETP (collected 23hrs after a recovery period (4xETP rec.), or five rounds of ETP-5xETP and transcript levels were compared to untreated, CON-neurons. Total RNA was isolated, and gene expression was accessed via RT-qPCR. **(C-D)** 4Cker, in the nearbait setting, was utilized to call significantly interacting regions in CON, 4xETP rec.-, 5xETP-, and 4xETP-NMDA-treated neurons across three 4C-seq replicates. Interactions, coverage plots, and TAD boundaries are as described in Figures 1A and 1B. **(E-F)** 4Cker, in the *cis* setting, was utilized to call significantly interacting regions in CON, 4xETP rec.-, 5xETP-, across three 4C-seq replicates. Interactions, coverage plots, and TAD boundaries are as described in Figures 1A and 1B. **(G)** Chromosomes 12 and 19 were segmented as described in Figure 1E. The number of regions with mapped 4C reads was assessed for CON-, 4xETP rec.-, 5xETP- and 4xETP-NMDA-treated neurons in each replicate. The average relative change in the number of contacted regions from the average CON level is shown for both the *Fos* (top) and *Npas4* (bottom) viewpoints. **(H)** The percent of intra- and inter-TAD reads (defined as described in Figure 1G) out of the total *cis* read amount were assessed for each replicate across all treatments. The pink symbols represent the intra-TAD values which are graphed on the left y-axis while the blue symbols represent inter- TAD values which are graphed on the right y-axis. This is shown for both the *Fos* (top) and *Npas4* (bottom) viewpoints.

Next, 4Cker was employed to assess the patterns of genome-wide significant contacts at both viewpoints in 4xETP rec., 5x ETP, and 4xETP-NMDA neurons. Analysis of nearbait interactions revealed that the interactome size at both viewpoints contracted substantially in 4xETP rec. neurons compared to untreated controls (Figure 3C and 3D). For instance, interactome size of the *Fos* and *Npas4* viewpoints were reduced by 32.3 % and 52.5 %, respectively (Figures 3C and 3D). Against this background, the *Fos* viewpoint interactome increased by only 15.1 % in 4xETP-NMDA-treated neurons and by only 13 % in 5xETP-treated neurons, compared to 4xETP rec. (Figure 3C). The nearbait interactome at the *Npas4* viewpoint increased by only 2.6 % in 4xETP-NMDA-treated neurons and contracted by 16.6 % in 5xETP-treated neurons, compared to 4xETP rec. neurons (Figure 3D). Meanwhile, *cis* interactomes contracted by 35.1% at the *Fos* viewpoint and by 59.1 % at the *Npas4* viewpoint in 4xETP rec. neurons relative to untreated controls and decreased further in 5xETP- and 4xETP-NMDA-treated neurons (Figures 3E and 3F). *cis* chromosome interactions at the *Fos* and *Npas4* viewpoints were reduced by 20.2 % and 22.2 %, respectively, in 4xETP-NMDA-treated neurons and by 14.3 % and 3.7 %, respectively, in 5xETP-treated neurons, compared to 4xETP rec. neurons (Figures 3E and 3F). Similarly, the number of discretely contacted regions were reduced substantially in 4xETP rec compared to untreated controls and were thereafter either unchanged or only modestly increased following subsequent DSB formation (Figures 3G). These reductions were distributed across all classes of chromatin states (Figures 3G). Furthermore, whereas singly ETP- and NMDA-treated neurons showed an increase in inter-TAD interactions, 4xETP rec, 5xETP, and 4xETP-NMDA showed a reduction in inter-TAD interactions and an increase in intra-TAD interactions (Figures 1G and 3H). Taken together, these results indicate that recurrent DSBs cause a remarkable pruning of ERG contacts with regions throughout the *cis* chromosome.

To determine the nature of chromosome interactions that do occur at ERG promoters, we expanded our analysis of significant contacts at *Fos* and *Npas4* viewpoints beyond their *cis* chromosomes. Analysis of the relative proportion of *cis*: *trans* mapped reads across all neuronal 4C-seq samples indicated that interactions of both viewpoints with regions on *trans* chromosome interactions were progressively increased with recurrent DSB formation (Figures 4A-4D). These observations suggest that potentiated ERG induction by recurrent DSBs correlates with a loss of *cis*, and a concomitant increase in *trans,* chromosome interactions with ERG promoters.

**Figure 4:**
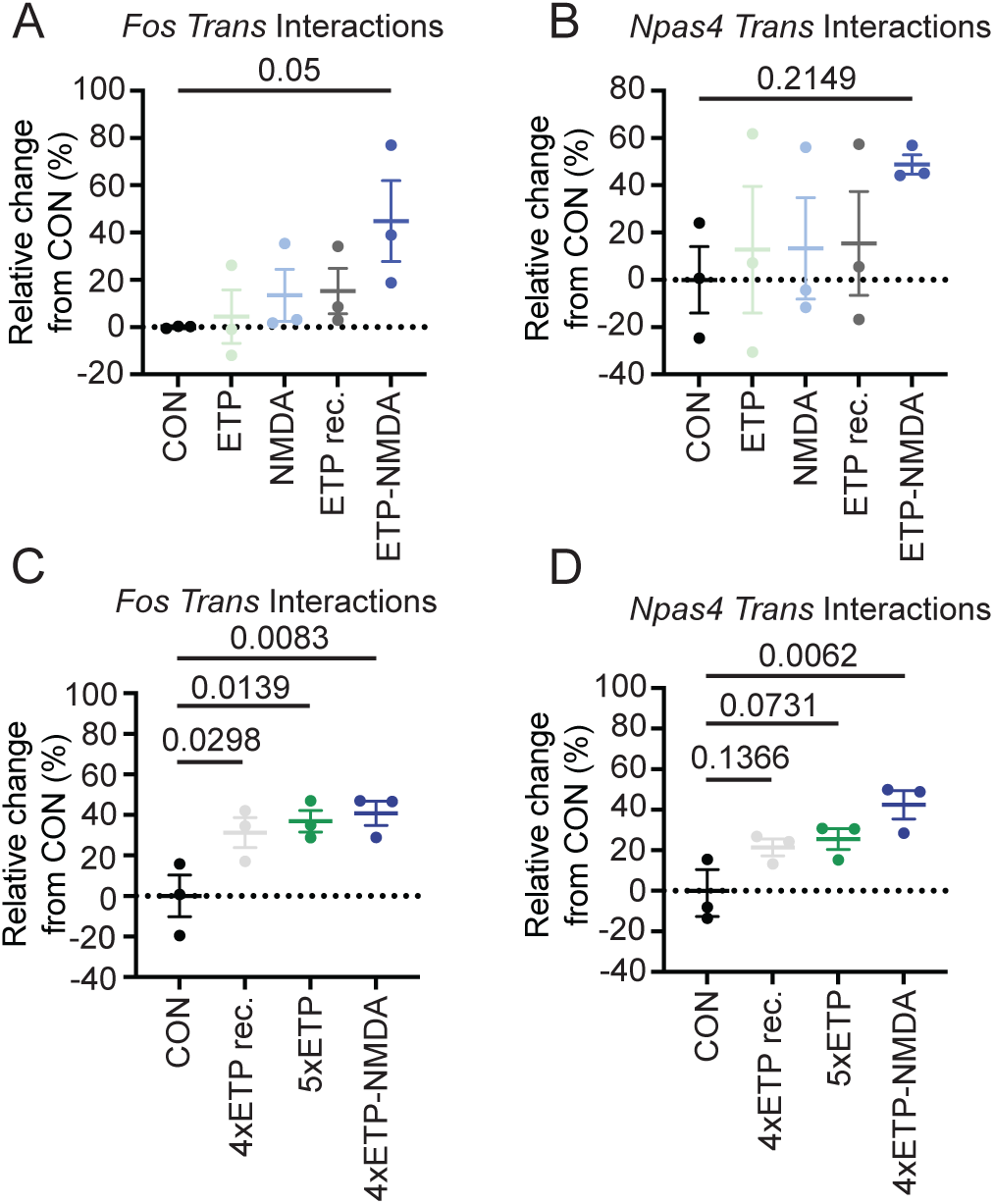
Recurrent TOP2B-mediated break formation is associated with increased *trans* interactions at the ERG promoters. **(A-B)** 4C reads were mapped to the genome and the ratio of *cis* to *trans* reads was assessed for each replicate of CON, ETP-, NMDA-, ETP rec.-, and ETP-NMDA-treated neurons for the *Fos* (A) or *Npas4* (B) viewpoint. The relative change in *cis: trans* ratio compared to CON neurons is shown for each treatment. **(C-D)** The ratio of *cis* to *trans* reads was assessed, as described in A, for each replicate of CON-, 4xETP rec.-, 5xETP-, and 4xETP-NMDA-treated neurons for the *Fos* (C) or *Npas4* (D) viewpoint. The relative change in *cis: trans* ratio compared to CON neurons is shown for each treatment.

### Recurrent targeted DSBs potentiate ERG transcription independently of DNA repair fidelity

The observation that activity-induced DSBs facilitate ERG transcription suggests that changes in DNA repair fidelity could also affect ERG transcription. For instance, it has been proposed that defective repair of recurrent activity-induced DSBs could lead to mutation accrual within ERG promoters and underlie their downregulation in the aging brain^7,25,35,36^. However, the effects of inaccurate repair following recurrent DSB formation on ERG transcription have not been assessed. To address this issue, we used the CRISPR-Cas9 system to generate targeted DSBs, which is known to induce indels at high frequency^47,48^. Additionally, we capitalized on findings that the transcription of ERGs, such as *Fos*, is also coupled to DSB formation in other systems and cell types. For instance, previous reports indicate that serum stimulation in HEK293T cells also induces TOP2B-mediated DSBs within the *FOS* promoter, which supports its rapid transcription^24^.

Based on these observations, HEK293T cells were transfected with Cas9 and one of two sgRNAs to target DSBs within regions 1 kb upstream of the *FOS* transcription start site (TSS) (Figure 5A). As a first step, the effects of Cas9-induced DSBs on *FOS* transcription and chromosome interactions at the *FOS* promoter were assessed 96 hours post-transfection. Cells transfected with sgRNAs targeted to the *FOS* promoter (CRISPR) showed about a 2-fold increase in *FOS* transcription compared to controls (CON) (Figure 5A). These results suggest that DSB formation within the *FOS* promoter is sufficient to upregulate its transcription in both proliferating and post-mitotic cells. Analysis of average normalized 4C-seq signals under these conditions revealed increased contacts throughout a 300 kb window surrounding the *FOS* viewpoint in CRISPR cells compared to CON (Figure 5B). Whereas contacts with putative enhancer elements was increased in CRISPR neurons (blue arrowheads), several regions with low chromatin accessibility also showed elevated interaction (orange arrowheads) (Figure 5B). Owing to the lack of available ChIP-seq datasets in HEK293T for several histone marks that were used to annotate chromatin states in neurons and the variability associated with using asynchronous cycling cells, two distinct strategies were used to assess contact frequencies across the *cis* chromosome with the *FOS* viewpoint: (1) human chromosome 14, which contains the *FOS* locus, was segmented into 500 bp-sized non-overlapping windows and the number of regions with mapped 4C contacts were compared; and (2) ChIP-seq peaks of select histone marks from available ENCODE data was used as a surrogate for functional states (promoters, actively transcribed gene bodies, enhancers, heterochromatin) and the number of regions with mapped contacts was compared^38^. Regardless of the strategy used, the number of contacted regions were increased in sgRNA-carrying CRISPR cells (CRISPR Gen 1) compared to CON Gen 1 cells (Figure 5C). For instance, on average, 3471 of the 500 bp *cis* chromosome windows showed contacts with the *FOS* viewpoint in CON Gen 1 cells, whereas this number increased by 18 % to 4101 regions in CRISPR Gen 1 cells (Figure 5C). Similarly, analysis across various histone marks indicated that increased contacts in CRISPR cells were distributed across functional states (Figure 5C). Thus, like with TOP2B-mediated DSBs in neurons, Cas9-induced DSBs at the *FOS* promoter in HEK293T cells were sufficient to increase its contacts with regions across the *cis* chromosome.

**Figure 5:**
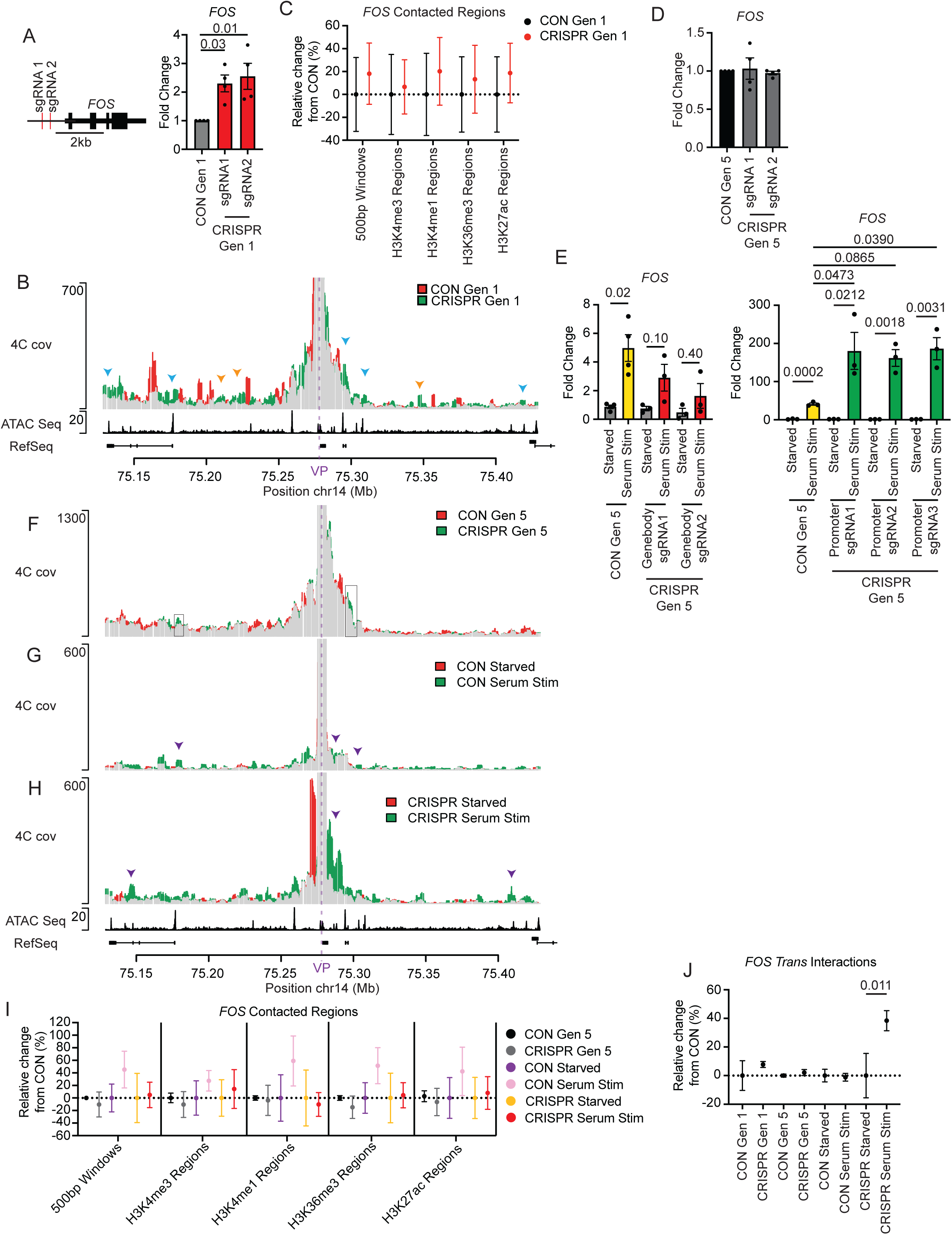
Recurrent targeted DSBs potentiate ERG transcription independently of DNA repair fidelity. **(A)** Diagram showing the location of the sgRNAs targeting 1kb upstream of the Fos TSS (left). *FOS* induction following transfection of CRISPR constructs (right). Gene expression was assessed via RT-qPCR of total RNA extracted from Hek-293T cells 96hrs after transfection with Cas9 and sgRNAs. **(B)** Overlap of the average, normalized 4C-seq signal between three replicates of CON Gen 1 and CRISPR Gen 1 in a 300kb region surrounding the *FOS* viewpoint (purple dashed line) is shown. The y-axis is the number of relative reads normalized to 1M *cis* reads. The x-axis is the position along chr14 in Mb and extends from 150kb upstream to 150kb downstream of the *FOS* promoter viewpoint. In track one the overlap (grey) between normalized 4C-seq signal in CON Gen 1 (red) and CRISPR Gen 1 (green) cells is shown. Blue arrow heads show regions of open chromatin with increased contact frequency in CRISPR Gen 1 cells. Orange arrow heads show regions of closed chromatin with increased contact frequency in CRISPR Gen 1 cells. Track two is previously published ATAC-Seq data from HEK293T to define regions of accessible chromatin, y-axis represents ATAC-seq signal. Track 3 is of hg38 RefSeq genes to help orient to the genomic location, thicker black bars represent protein coding regions, while narrow lines correspond to intronic regions. **(C)** Chromosome 14 was split into non-overlapping 500bp windows and was split into functionally distinct regions with available histone mark ChIP-seq peak calling data. The number of regions with mapped 4C reads was assessed for each set of regions in each replicate in CON Gen 1 and CRISPR Gen 1 HEK293T cells. The average relative change in the number of contacted regions from the average CON Gen 1 level is shown for the *FOS* viewpoint. **(D)** HEK293T cells were transiently transfected with sgRNA guides targeting the *FOS* promoter. Generation 1 began 96 hrs post transfection and both CON and CRISPR cells were passaged for five generations (Gen 5). Total RNA was isolated from HEK293T cells, and gene expression was accessed via RT-qPCR. **(E)** HEK293T cells were transiently transfected with guides targeting either exons (left) or the promoter (right) of *FOS*. Generation 1 was 96 hrs after transfection and both CON and CRISPR cells were passaged for five generations as shown in Figure S5A and then subjected to serum starvation for ∼16.5 hrs followed by serum stimulation for 30 min. Total RNA was extracted, and gene expression was accessed via RT-qPCR. **(F)** Overlap (grey) of the average normalized 4C-seq signal between CON Gen 5 (red) and CRISPR Gen 5 (green) conditions in a 300kb region surrounding the *FOS* viewpoint (purple dashed line) is shown. The y-axis is the number of relative reads normalized to 1M *cis* reads and the x-axis is the position along chr14 in Mb. The black boxes represent the regions that showed increased contact in CRISPR Gen 1 cells that still have increased contact five generations later. **(G)** The overlap (grey) between normalized 4C-seq signal in CON (Gen 5) Starved (red) and CON (Gen 5) Serum Stimulated (green) cells is shown. The y-axis is the number of relative reads normalized to 1M *cis* reads and the x-axis is the position along chr14 in Mb. Arrowheads show putative enhancer regions with stimulus-induced contact frequency increases. **(H)** The overlap (grey) between normalized 4C-seq signal in CRISPR (Gen 5) Starved (red) and CRISPR (Gen 5) Serum Stimulated (green) cells is shown. Arrowheads show putative enhancer regions that gain increased contact following serum stimulation in CRISPR Gen 5 cells versus CON Gen 5 cells. ATAC-Seq and RefSeq tracks are the same as provided in B. **(I)** Chromosome 14 was segmented as described in C. The number of regions with mapped 4C reads was assessed for each replicate in CON/CRISPR Gen 5, CON Serum starved/stimulated, and CRISPR Serum starved/stimulated HEK293T cells. The average relative change in the number of contacted regions from the average timepoint matched CON sample is shown for the *FOS* viewpoint. **(J)** 4C reads were mapped to the genome and the ratio of *cis* to *trans* reads was assessed for each replicate of all HEK293T *FOS* 4C-seq samples. The relative change in *cis*:*trans* ratio compared to time point-matched CON HEK293T cells is shown.

Whereas the fidelity with which activity-induced DSBs are repaired remains unclear, Cas9-induced DSBs are often repaired using error-prone repair pathways^47,48^. We reasoned therefore that analyses following Cas9-induced DSB could allow us to assess the impact of low-fidelity DSB repair on ERG transcription. To this end, HEK293T cells were again transiently transfected with Cas9 and sgRNAs targeting the promoter of *FOS*. Cells were then allowed to recover and were passaged up to 5 times (Generation 5, Gen 5) (Figure S5A). Immunofluorescence and western blotting experiments confirmed the expression of Cas9 following transfection and its gradual loss as a function of passage number (Figures S5A-S5C). Previous reports indicate that continuous Cas9 expression in this manner can result in multiple rounds of cutting and repair until indels that prevent target recognition by the sgRNA are generated^47,49^. Following the loss of Cas9 (CON Gen 5 and CRISPR Gen 5), cells were serum starved for ∼ 16.5 hours and then stimulated with 18 % fetal bovine serum (FBS) for 30 min. The effects of prior cycle(s) of Cas9-mediated DSB formation and erroneous repair on stimulus-driven *FOS* transcription and chromosome interactions with the *FOS* promoter were then assessed.

*FOS* expression in CRISPR Gen 5 cells was indistinct from CON Gen 5 cells (Figure 5D), suggesting that recurrent DSB formation did not alter baseline levels of *FOS* expression. Serum stimulation of control cells caused a robust induction of *FOS*, as indicated by qRT-PCR experiments (Figure 5E). Additionally, and consistent with expected effects of indels, serum-induced *FOS* transcription was reduced in HEK293T cells transfected with Cas9 and sgRNAs that targeted DSBs within the exons of *FOS* (Figure 5E). By contrast, serum-induced *FOS* transcription was significantly potentiated in HEK293T cells transfected with Cas9 and sgRNAs directed to the *FOS* promoter (CRISPR Gen 5 cells) compared to controls (Figure 5E). These results indicate that prior DSB formation within ERG promoters can potentiate ERG transcription even when such DSBs are repaired inaccurately.

To determine how potentiation of *FOS* transcription by Cas9-induced DSBs relates to chromatin interactions at the *FOS* promoter, 4C-seq was performed in CON and CRISPR Gen 5 cells following serum starvation and serum stimulation. Comparison of local 4C signals indicated a modest reduction in CRISPR Gen 5. cells compared to CON Gen 5 cells in many areas, yet contacts with several regions that were increased immediately following CRISPR were also maintained (Figures 5B and 5F). Serum stimulation following CRISPR rec. again stimulated contacts throughout the 300 kb window (Figure 5H). Overall, local contact profiles in unstimulated and serum-stimulated CRISPR rec. cells were reminiscent of those in ETP rec. and ETP-NMDA neurons (Figure S3A, S3F, 5F-5H). To further examine this possibility, *cis* chromosome contacts with *FOS* promoter were compared in CON and CRISPR Gen 5. cells as a function of serum stimulation, as described above. Analysis of mapped interactions with 500bp non-overlapping segments of human chromosome 14 indicated a marked increase in contacts following serum stimulation in CON cells and this increase was distributed across regions marking promoters, enhancers, and gene bodies (Figure 5I). Whereas contacts with *cis* chromosome regions were also elevated in CRISPR Gen 5 cells following serum stimulation, this increase was less pronounced compared to CON Gen 5 cells that were serum starved and stimulated (Figure 5J). For instance, whereas serum stimulation caused an ∼ 45 % increase in the number of 500 bp segments in chromosome 14 contacted by the *FOS* promoter in CON Gen 5 cells, but only an ∼ 4 % increase in such contacts in CRISPR Gen 5 cells (Figure 5I). In contrast to a modest increase in *cis* chromosome contacts, serum stimulation in CRISPR Gen 5 cells caused a ∼ 38 % increase in *trans* chromosome contacts, while *trans* chromosome contacts in serum-stimulated CON Gen 5 cells, CON/CRISPR Gen 5 cells, and CON/CRISPR Gen 1 cells remained unaffected (Figure 5J). Overall, these results indicate that recurrent DSBs can reconfigure chromosome contacts and potentiate ERG transcription even under conditions of low DNA repair fidelity.

## DISCUSSION

In this study, we attempted to assess how neuronal activity-induced DSBs impact chromosome interactions at ERG promoters and ERG transcription. Using multiple strategies to generate DSBs within the promoters of the ERGs, *Fos* and *Npas4*, and performing 4C-seq, we show that DSBs dynamically reconfigure chromatin contacts at ERG promoters. Concurrently, our results shed new light on the links between 3D genome organization and ERG transcription.

Previous reports suggest that activity-induced *de novo* loops formed between ERG promoters and specific enhancers are crucial for ERG transcription^11–13^. Although enhancer interactions were also detected in our 4C-seq experiments, we find no evidence for directed promoter-enhancer looping at ERGs. Instead, neuronal stimulation expanded the interactome of ERG promoters with diverse chromatin states, with enhancers comprising only a small fraction of such contacts. Whereas the significance of interactions between ERG promoters and regions other than enhancers is unclear, several studies indicate that regions lacking canonical chromatin signatures of enhancers could still possess enhancer activity^50–52^. Encounters between such regions and ERG promoters could thus boost ERG transcription. These results suggest that activity-induced contacts with ERG promoters could initially form through increased mobility-type mechanisms rather than by their directed looping with specific enhancers^53,54^. Such a scenario is consistent with observations that promoter-enhancer interactions at neuronal ERGs are unaffected by the loss of cohesin^21^. Neuronal activity could increase chromatin mobility in several ways; however, we propose that activity-induced DSBs are an important contributor to chromatin dynamics at ERG promoters. Signaling activated by DSB formation is known to alter the localization of DSB sites and the biophysical properties of chromatin fibers, such as their compaction and stiffness, in a context-dependent manner^55–58^. The mobility gained from these events could explain how DSB formation is sufficient to enlarge the interactome of ERG promoters.

Notably, the inducibility of ERGs did not scale with the interactome size of ERG promoters and a single round of DSB formation caused only a modest upregulation of ERGs compared to their induction following neuronal stimulation. Separate from triggering DSBs within ERG promoters, neuronal activity-dependent signaling also recruits chromatin remodelers, histone acetyltransferases, and transcription factors to the *cis*-regulatory elements of ERGs^7^. The actions of these factors could either elevate chromatin mobility or support ERG transcription independently of 3D chromatin organization. For instance, promoter H3K27ac alone accounted for a significant proportion of the variance in neuronal activity-dependent transcription^12^. Together, these observations suggest how DSB and neuronal activity-dependent signaling cooperatively drive the rapid and robust transcriptional induction of ERGs.

We report here that ERGs with a history of incurring TOP2B-mediated DSBs within their promoters show higher transcriptional induction in response to subsequent neuronal stimulation. Experiments in HEK293T cells indicate that the ability of DSBs to potentiate ERG transcription is heritable. These observations are reminiscent of the phenomenon of transcriptional memory and further highlight the regulatory impact of DSB formation on ERG transcription^59^. How DSBs prime higher ERG induction in subsequent rounds of stimulation is unclear. Whereas current models invoke certain histone marks and transcription factors as carriers of transcriptional memory, chromatin reorganization could also prime gene activity states^59^. ERG promoter contacts and accessibility changes that are retained long after the cessation of ERG transcription and DSB repair could establish a new ground state that allows for higher ERG induction in response to an ensuing stimulus^11,12,60,61^. By contrast, the remarkable loss of *cis* chromosome contacts with ERG promoters caused by recurrent DSBs could also potentiate ERG transcription by culling interactions with repressive *cis* regulatory elements^62^. Such a model would be analogous to those in which transcriptional memory is attained by the loss of either DNA methylation or repressive histone methylation^59,63,64^. Notably, the loss of *cis* chromosome contacts following recurrent DSB formation was accompanied by elevated *trans* interactions. While the significance of increased *trans* chromosome interactions remains ambiguous, DSB-containing domains were recently shown to self-segregate from other chromatin regions to form a distinct chromatin compartment (the D compartments)^65^. The D compartment also showed elevated *trans* interactions, recruited damage-responsive genes, and activated their transcription^65^. Future imaging-based approaches should elaborate whether ERGs become similarly compartmentalized, and whether such a mechanism underlies their potentiated induction.

While activity-induced DSBs regulate ERG transcription, their formation carries the risk of mutation and genomic instability. Unexpectedly, we observe that ERGs retain robust inducibility despite accumulating mutations within their promoters. This resilience suggests that the risk posed by mutations in ERG promoters for gene expression could be relatively low under normal conditions. Whether such risks escalate during aging and age-related disorders warrants further investigation^36^. Whereas mutational accrual may not affect ERG inducibility, delayed and defective DSB repair could dysregulate either the kinetics or the duration of ERG transcription, and compromise neuronal functions^22^. Future efforts aimed at understanding how activity-induced DSBs are repaired and assessing the effects of defective DSB repair *in vivo* should address this issue.

## ACKNOWLEDGMENTS

This work was supported by funding from the NIMH R01MH120132 and R01MH130399 to R.M., and by NIH T32GM109776 and T32HL139438 to L.H.

## AUTHOR CONTRIBUTIONS

Methodology – L.H. R.M.; Investigation – all authors; Writing – L.H., R.M.; Funding Acquisition – R.M., L.H.; Resources – R.M.; Supervision – R.M.

## RESOURCE AVAILABILITY

### Lead Contact

Further information and requests for resources and reagents should be directed to and will be fulfilled by the lead contact, Ram Madabhushi.

### Materials Availability

This study did not generate new unique reagents.

### Data availability

All sequencing data generated in this study has been deposited at GEO and will be publicly available following peer-review and publication. The existing, publicly available data that was analyzed for this study are listed in the Supplementary Table 1 along with accession numbers.

## METHODS

### EXPERIMENTAL MODEL DETAILS

#### Cell Culture and Treatments

Dissociated primary cortical neurons were cultured as previously described^22,31,40,66^ and supplemented with L-glutamine, penicillin/streptomycin, and B27. All ETP treatments were conducted with 10μM final concentration, for at most 1hr. NMDA treatments were with 50μM and lasted for 10 min followed by a recovery in conditional neuron media. Figure S2A and Figure S4A outline the recurrent treatment paradigms used in this study.

Human embryonic kidney cells, HEK293, were maintained in Dulbecco’s modified Eagle’s medium (DMEM, high glucose) supplemented with 10% FBS and 5% penicillin/streptomycin. Serum starvation of the Hek293 cells was done in 0.5% FBS for ∼16.5hrs and then serum stimulated in 18% FBS as described previously^24^.

### METHOD DETAILS

#### 4C-seq

4C-seq was performed as described previously^34^. Approximately 10 million primary cortical neurons (DIV13) or HEK293T cells were cross-linked in 2% formaldehyde for 10 min at room temperature and quenched with glycine (0.13M final concentration). Cells were incubated for 20 min on ice in lysis buffer (50 mM Tris-HCl pH 75, 150 mM NaCl, 5 mM EDTA, 0.5% NP-40, 1% TX-100, and protease inhibitors). Nuclei were resuspended in 450 μl 1.2x CutSmart buffer and permeabilized with 15 μl 10% SDS for 1hr and 75 μl 20% Triton X-100. A sample of undigested chromatin was collected to assess chromatin integrity. DNA was digested with 100U NlaIII at 37°C for 3 hrs while shaking at 750 rpm and then an additional 100 U of NlaIII was added and incubated overnight. The extent of chromatin digestion was assessed via gel electrophoresis. NlaIII was deactivated by incubating samples for 20 min at 65°C. Chromatin was diluted to 7 ml, supplemented with 10X ligation buffer, and 4000 U NEB T4 DNA Ligase for overnight ligation at 16°C while shaking. Ligation efficiency was checked by gel electrophoresis. Crosslinks were reversed and proteins digested with proteinase K (30 μl of 10 mg/ml) overnight at 65°C. DNA was extracted via phenol:chloroform and precipitated with ethanol. DNA was then digested overnight at 37°C with 50 U DpnII supplemented with 10X DpnII Buffer while shaking. Second digestion efficiency was determined via gel electrophoresis. DpnII was deactivated at 65°C for 20 min. DNA circularized in a 7 ml reaction, supplemented with 10X ligation buffer, and 4000 U NEB T4 DNA Ligase overnight at 16°C with shaking. Indexed Illumina sequencing adapters were then introduced to ligation fragments of interest with a two-step PCR strategy using Expand Long Template Polymerase starting with 200 ng of 4C library template. Viewpoint-specific primers are listed in Supplementary Table 2. The first round of amplification was performed with the following thermocycler conditions: 94°C for 2 min; 94°C for 10 sec, Viewpoint specific temperature (*Fos*-68°C, *Npas4*-57°C, *FOS*-56°C), for 1 min, 68°C for 3 min, for 16 cycles; and 68°C for 5 min. Five different PCR reactions per sample were pooled and 50 μl of the pool was used to perform 0.8 x AMPure XP bead purification. 5 μl of purified round 1 PCR product was then used for the second round of amplification conducted with the following cycling conditions: 94°C for 2 min; 94°C for 10 sec, 60°C for 1 min, 68°C for 3 min for 20 cycles; and 68°C for 5 min. PCR clean-up was done using Qiagen PCR Purification kit. DNA fragment size of final, clean PCR products was assessed via Tapestation (Agilent). Library concentrations for each sample was assessed using fluorometry (dsDNA high sensitivity Qubit assay, Thermo) and RT-qPCR (KAPA Biosystems) library quantification. These were used to pool equimolar 4C-seq libraries of all samples and replicates. Experiments were conducted in triplicates and in parallel with RT-qPCR to capture the associated transcriptional state. Libraries were sequenced on an Illumina NextSeq 500 instrument using V2.5 reagents to a depth of at least 1 million usable mapped *cis* reads per sample.

#### 4C-seq Analysis

Processing of 4C-seq data was performed using the 4Cker pipeline^41^ (interactome calling) and the pipe4C pipeline^67^ (visualization and BAM file generation). 4Cker mapping was performed using Bowtie2 to a reduced genome consisting of all unique x-nt (nucleotide count differs depending on sequencing run and viewpoint specific primers) regions surrounding DpnII sites from the hg38 or mm10 reference genome allowing for zero mismatches. To define regions with significant interaction with the viewpoint we used either *cisAnalysis* (*cis*) or *nearBaitAnalysis* (nearbait) of 4Cker inputting three replicates per sample using the indicated k-values (Supplementary Table 3): CON, ETP, NMDA, ETP rec. ETP NMDA – Fos nb k=4 cis k=12; Npas4 nb k=3, cis k=11 CON, ETP x 4 rec., ETP x 5, ETP x 4 NMDA – Fos nb k=7 cis k=15; Npas4 nb k=4, cis k=5 The BED file outputs from both nb and cis were intersected with chromosome states using bedtools *intersect* and then bedtools^68,69^ *genomecov* to calculate the amount of coverage (in bp) of each chromatin state in each sample’s interactome. Relative change in interactome size from untreated, CON neurons are shown.

Mapping with pipe4C was done with the following parameters: normalization to 1 million reads in cis, non-blind fragments only, window size 21, top 2 read counts removed. Profile overlays were produced using R and the .rds files output from pipe4C. BAM files were also generated by pipe4C which were used to determine how many of the discrete regions had mapped contacts in them. Regions on mm10 chr12 and chr19 were segmented by ChromHMM states in neurons and hg37 chr14 was segmented with histone mark ChIP-Seq called peaks or with 500bp tiled windows in Hek293T cells. These defined regions were used with bedtools *multicov* along with the individual BAM files to assess how many unique regions were contacted in each sample. The number of contacted regions were compared to the average control of each replicate to determine the relative change from control which is shown. Bedtools multicov was also utilized with compartmental (to calculate the percent of reads in each compartment on mm10 chr 12 and chr19), chromosomal (used to calculate *cis*/*trans* read ratios), and TAD (used to calculate intra-TAD and inter-TAD contact percentage on the *cis* chromosome) beds to calculate the number of mapped reads in each replicate.

#### Chromatin State Annotation

Annotation of chromatin states was done using ChromHMM^37,70^ as described previously^40^. BED files corresponding to each chromatin state were used to examine the number or discrete regions that are contacted by the promoters. The 15 chromatin states were reclassified into 4 broad functional groups: enhancer, promoter, heterochromatin, and genebody, see figure S1D for clustering data.

#### Hi-C analysis

Previously published Hi-C data from mouse cortical neurons was used to identify chromosome compartments using the Juicer^46^ suite. Chromosome compartments were called using *eigenvector* command with a 500kb resolution and KR normalization. TADs were called using Arrowhead with 5 kb resolution and KR normalization.

#### RT-qPCR

Primary cortical neurons or HEK293T cells were plated in 24 well plates and total RNA was extracted with TRIzol reagent (Invitrogen) following the manufacturer’s instructions and precipitated with ethanol. cDNA was synthesized from 1ug of total RNA using an RT-PCR EcoDry Premix (Clontech). RT-PCR reactions were performed using SssAdvanced Universal SYBR Green Supermix using the indicated primers (Supplementary Table 2). Data are shown as fold change relative to unstimulated controls. The number of dots per graph represent the n for every experiment, and p-values < 0.05 for comparisons are indicated on the graphs. The data were analyzed by one-way ANOVA as appropriate using Tukey’s post-hoc tests.

#### Western blotting

Cells were plated in 6 well plates and lysed in RIPA buffer (50mM Tris-HCl, pH 7.5, 150mM NaCl, 1mM EDTA, 0.1% SDS, 1% IGEPAL-CA630, 1% sodium deoxycholate, protease and phosphatase inhibitor cocktails) and prepared for western blotting as described previously^66^. Membranes were incubated overnight with primary antibodies: γH2AX (MilliporeSigma, 05-636, (RRID: AB_309864)), H2AX (Abcam, ab11175, (RRID: AB_297814)), β-actin (Bio Rad, VMA00048, (RRID: AB_2923317)), and Cas9 (Abcam, ab191468). Thereafter the membranes were washed in TBS-T, and incubated with 1hr with goat anti mouse or rabbit IR dye-conjugated secondary antibodies (LI-COR, 92668170, (RRID:AB_10956589) and 82708365, (RRID:AB_10796098)). Graphs show normalized optical density and expressed as fold change over unstimulated controls. Each dot in the bar graph represents the result of an independent experiment/biological replicate performed on different days using neurons cultured from independent mouse litters.

#### Immunocytochemistry

Cells plated on coverslips were fixed with 4% paraformaldehyde for 10 min, washed with PBS and incubated with blocking buffer (5% normal goat serum, 0.3% Triton X-100, in PBS) at room temperature overnight. Then, the cells were incubated with anti-GFP antibody (1:1000 dilution, AVES GFP-1010) in blocking buffer overnight at 4°C. The following day the cells were incubated with secondary antibody goat anti-chicken IgG-Alexa Fluor 488 (Thermo Fisher, A11039) in blocking buffer at room temperature for 1hr and washed. DNA was stained with Hoechst 33342 for 5 min and washed before being mounted with ProLong Diamond Antifade (Thermo Fisher). All images were acquired using a Zeiss LMS800 confocal.

#### Cell Titer-Glo

Cells were cultured in 24 well plates, stimulated with the indicated conditions, and allowed to recover for 8hrs post stimulation. Cells were lysed according to CellTiter-Glo (Promega) manual specifications. Cells were incubated for 10 min with agitation. Each well had three technical replicates run along with each sample having four biological replicates. Luminescence levels were assessed with CLARIOstar plate reader. The change in average luminescence from the control cells was assessed.

#### Propidium Iodide Staining

Cells were cultured in 24 well plates, stimulated with the indicated conditions, and allowed to recover for 8hrs post stimulation. Propidium iodide (PI) was added to the culture media to a final concentration of 50μg/mL. Cells were allowed to incubate at 37°C for 15 min. PI staining levels were checked via matrix well scanning on the CLARIOstar plate reader at 535nm excitation. Triton X-100 was added to a final concentration of 0.1% to the culture media to permeabilize all cells to get a maximal PI labeling amount per well. Cells were incubated with Triton X-100 for 15 min at 37°C. PI levels were checked again using the matrix well scanning settings. We then calculated each well’s percentage of total death (PI staining post Triton X-100). Deviations from the control level of death was assessed and plotted. Representative whole well scans are also shown. Each data point is representative of n for the experiment, one-way ANOVA’s were performed to determine if there was a treatment effect on the membrane integrity.

#### Chromatin Immunoprecipitation

©H2AX-ChIP was performed for 3-4 biological replicates essentially as described previously^31,40,66^. Samples were then incubated with 7 μg of anti-©H2AX (Abcam -ab2893). Final DNA ChIP and Input samples were then subjected to quantitative PCR using the indicated primers (Supplementary Table 2). Data are shown as fold change relative to unstimulated controls. One-way ANOVAs with Tukey’s post-hoc test were conducted to test for statistical significance.

## SUPPLEMENTARY FIGUR LEGENDS

**Figure S1:**
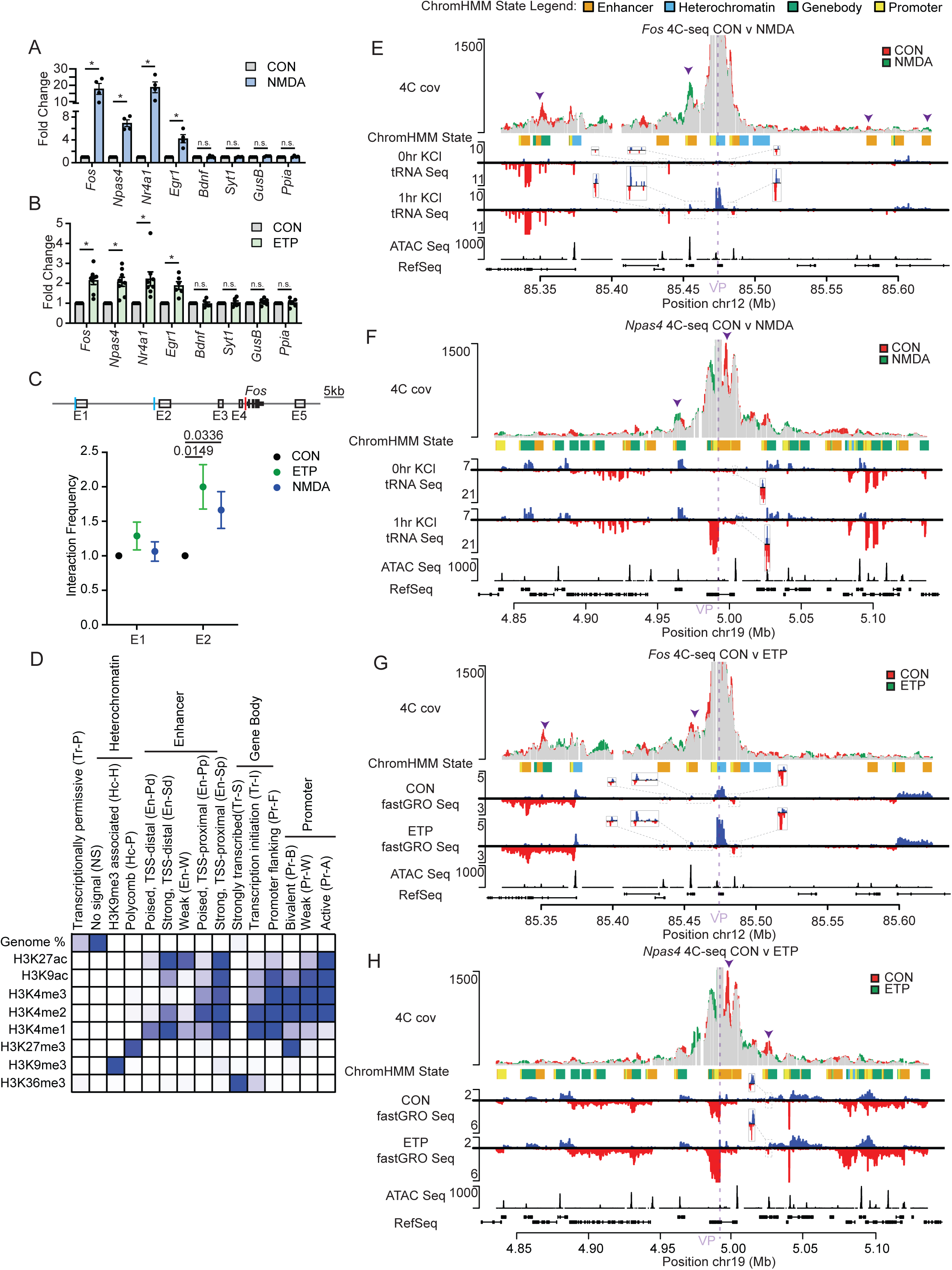
TOP2B mediated break formation induces ERGs and alters local contact frequencies. **(A)** DIV13 primary cortical neurons were treated with NMDA (50μM, 10 min followed by washout and a 20 min recovery). Total RNA was extracted, and gene expression was accessed via RT-qPCR. **(B)** DIV13 primary cortical neurons were treated with ETP (10μM, 30 min). Total RNA was extracted, and gene expression was accessed via RT-qPCR. **(C)** Top: Diagram of previously characterized activity-dependent Fos enhancers E1-E5 along with the location of the bait primer (red) in the promoter and interaction probes (blue) directed at E1 and E2. Bottom: DIV13 primary cortical neurons were stimulated with either NMDA (50 μM, 10 min followed by washout and a 50 min recovery) or ETP (10 μM, 30 min). Cells were subsequently cross-linked and nuclei lysed. Chromatin was then digested with BgIII and SacI overnight followed by ligation and DNA purification. The purified DNA was used as a qPCR template with the primers indicated and interaction frequencies at both E1 and E2 were calculated relative to untreated, CON neurons. **(D)** Eight available ChIP-seq datasets of histone marks in P0 forebrain tissue on ENCODE. ChromHMM was utilized to build a genome segmentation model into 15 functionally distinct chromatin states. A heat map is shown with each state and the relative enrichment of each histone mark and total genome percentage. The 15 chromatin states were grouped into broader functional classes: heterochromatin, enhancer, gene body, and promoter regions. **(E-F)** Overlap (grey) of the normalized 4C-seq signal between untreated CON-(red) neurons and NMDA-treated (green) neurons in a 300kb region surrounding the *Fos* (E) or the *Npas4* (F) viewpoint is shown. y-axis is the number of relative reads normalized to 1M *cis* reads (Track 1). ChromHMM states of local regions are shown based on the four broad functional types of regions: enhancers (orange), heterochromatin (blue), gene body (green), promoter (yellow) (Track 2). To map activity dependent enhancers, previously published^9^ total RNA seq reads in primary cortical neurons either 0hr (Track 3) or 1hr (Track 4) after KCl stimulation were remapped to mm10 and stranded bigwigs were generated and are shown surrounding the viewpoints. The blue signal is the forward strand and the red signal is the reverse strand. Zoom-in views are shown of annotated enhancers that have stimulus specific increases in transcription. To visualize regions of open chromatin we utilized available ATAC-Seq from ENCODE, bigwig signal is shown (Track 5). To orient to the location in relation to other genes, RefSeq genes are shown where the thicker boxes represent protein coding regions, and the thin bar represents introns (Track 6). Arrow heads highlight regions of interest that incur changes in contact frequency. **(G-H)** Overlap (grey) of the normalized 4C-seq signal between untreated CON- (red) neurons and ETP-treated (green) neurons in a 300kb region surrounding the *Fos* (G) or the *Npas4* (H) viewpoint is shown. Y-axis is the number of relative reads normalized to 1M *cis* reads (Track 1). ChromHMM states of local regions are shown based on the four broad functional types of regions: enhancers (orange), heterochromatin (blue), gene body (green), promoter (yellow) (Track 2). To map enhancers that are activated by ETP treatment, previously published^40^ fastGRO-seq in primary cortical neurons either control (Track 3) or ETP treated (Track 4) is shown via stranded bigwigs. The blue signal is the forward strand and the red signal is the reverse strand. Zoom-in views are shown of annotated enhancers that have stimulus specific increases in transcription. To visualize regions of open chromatin we utilized available ATAC-Seq from ENCODE, bigwig signal is shown (Track 5). To orient to the location in relation to other genes, RefSeq genes are shown where the thicker boxes represent protein coding regions, and the thin bar represents introns (Track 6). Arrow heads highlight regions of interest that incur changes in contact frequency.

**Figure S2:**
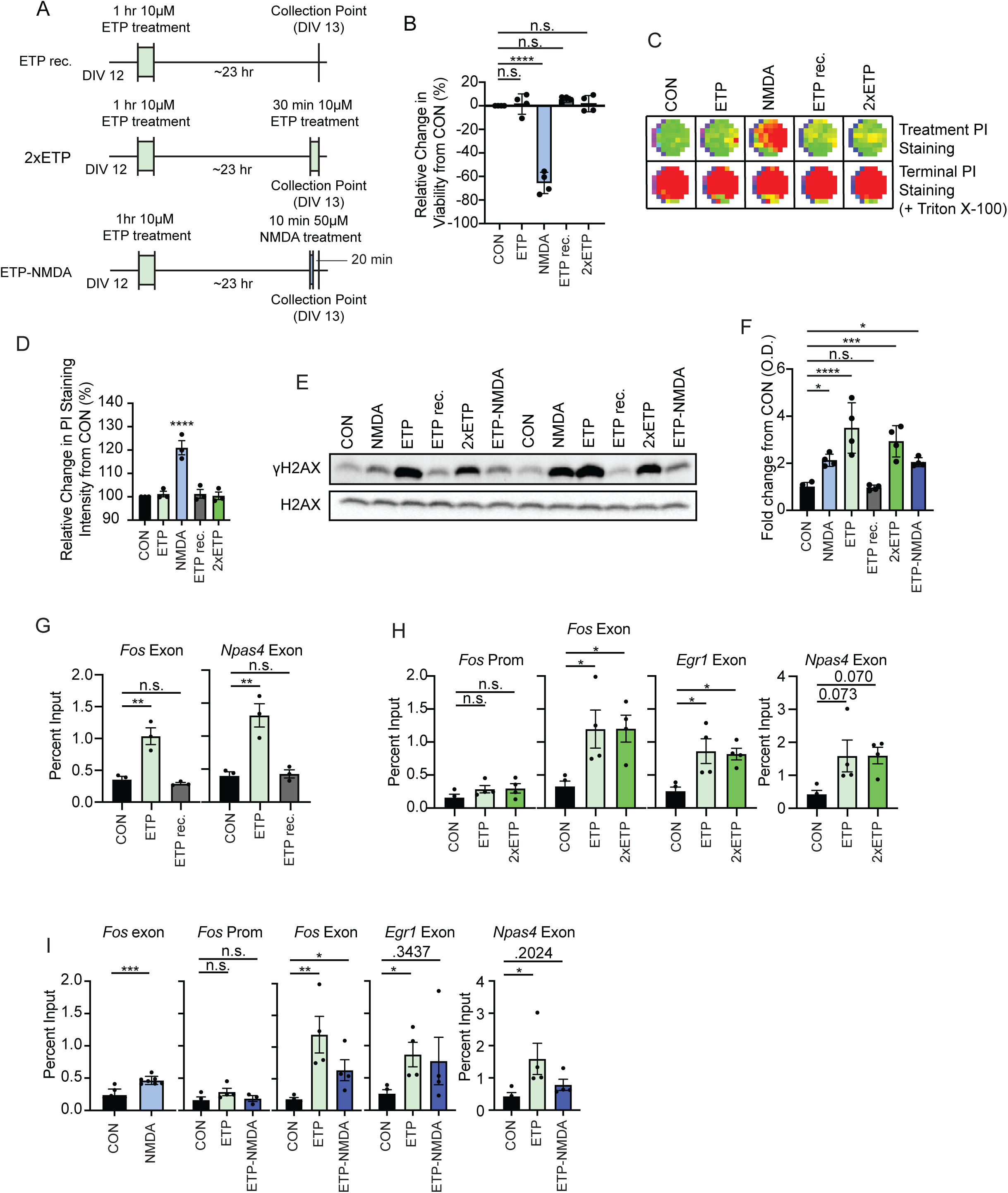
Repeated TOP2B-mediated break formation can occur in the ERG promoters. **(A)** Schematic diagram of the ETP rec., 2xETP, and ETP-NMDA treatments. DIV12 neurons are stimulated with ETP (10M, for 1hr) followed by a ∼23hr recovery period. The neurons are either collected after the recovery period (ETP rec.) or a second round of break formation is triggered by either a second ETP treatment (10M, 30min) (2xETP) or an NMDA treatment (50M, 10 min stimulation with a washout and a 20 min recovery period) (ETP-NMDA). **(B)** ATP levels were quantified as a proxy for cellular health. Neurons were subjected to the indicated treatments. Cells were allowed to recover for 8hrs after the final stimulation or after the 23hr recovery period and were then lysed and ATP levels quantified by the CellTiter-Glo (Promega) kit according to the manufactures specification. Average ATP levels were calculated across replicates and the relative change in ATP levels from CON neurons was determined and is shown as a percent. **(C-D)** Membrane integrity was assessed via propidium iodide (PI) staining. Neurons were subjected to the indicated treatments. Cells were allowed to recover for 8hrs after the final stimulation or after the 23hr recovery period. Cells were then incubated with PI that stain the cells with compromised membranes. PI staining was assessed via well scanning and representative heatmaps are shown (C) for each treatment. Cells were then treated with Triton X-100 and PI staining levels was assessed again to quantify the maximum PI staining levels of each well, Terminal PI Staining (C). Average treatment PI staining was normalized for each well based on the terminal PI staining values. The relative change in normalized PI staining intensity from CON neurons is shown (D). **(E-F)** Neurons were subjected to the indicated treatments. Neurons were collected and lysed 20 min after the final treatment or the 23hr recovery period. Western blots of total H2AX and ãH2AX for two representative replicates are shown (E). ãH2AX levels were normalized to total H2AX levels and are quantified for 4 replicates and shown (F). **(G)** Neurons were treated with ETP for 1hr and collected either 0hr (ETP) or 24hrs (ETP rec.) after stimulation. ©H2AX levels were assessed via ChIP-qPCR using primers targeting exons in *Fos* and *Npas4*. The percent input of each ChIP sample is shown for each replicate. **(H)** Neurons were treated with either ETP or 2xETP and were collected 20 min after the final stimulation. ©H2AX levels were assessed via ChIP-qPCR using primers targeting the *Fos* promoter, and exons in *Fos*, *Npas4, Egr1*. The percent input of each ChIP sample is shown for each replicate. **(I)** Neurons were treated with either ETP or ETP-NMDA and were collected 20 min after the final stimulation. ©H2AX levels were assessed via ChIP-qPCR using primers targeting the *Fos* promoter, and exons in *Fos*, *Npas4, Egr1*. The percent input of each ChIP sample is shown for each replicate.

**Figure S3:**
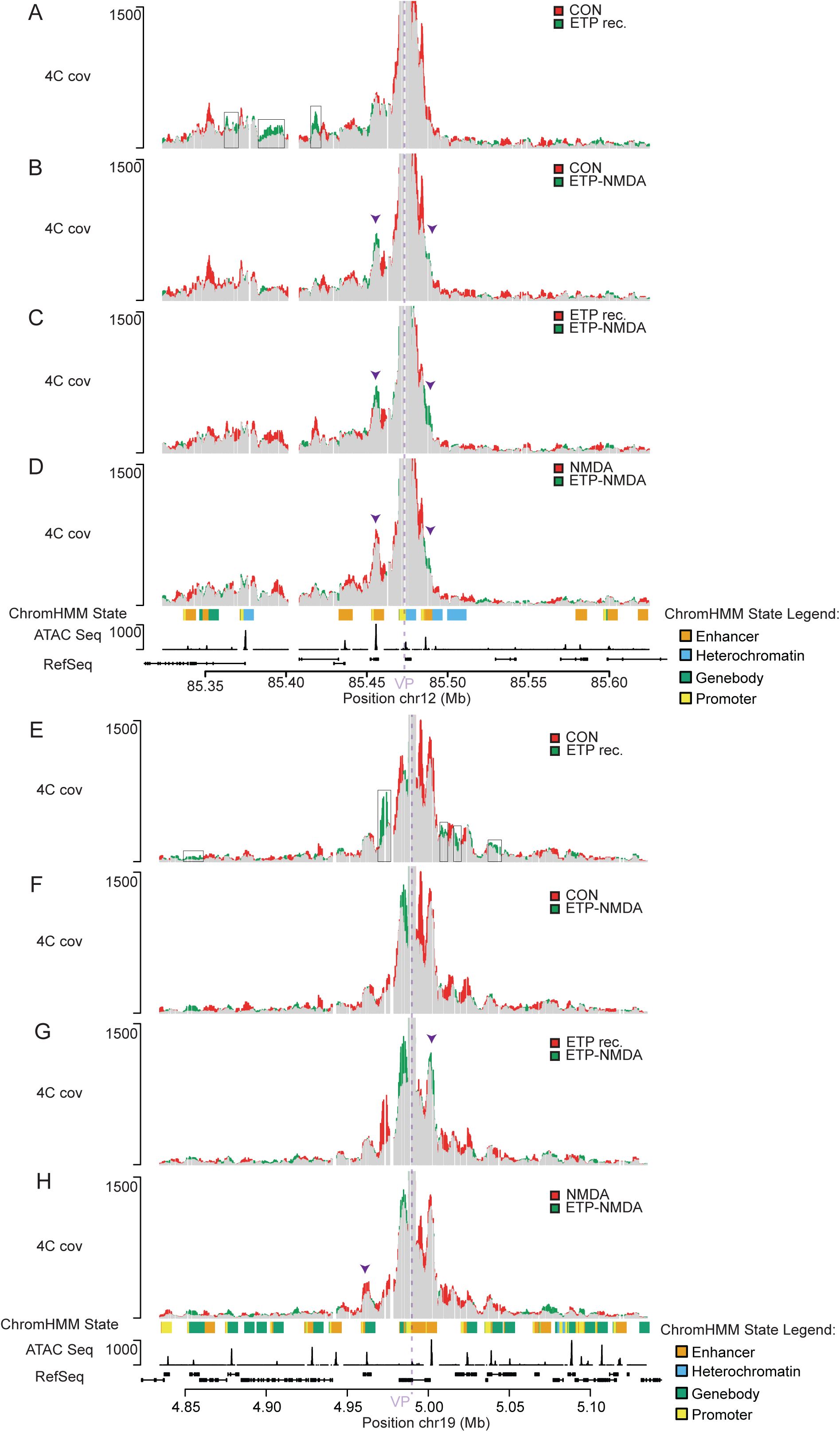
A prior TOP2B-mediated break formation subsequent to neuronal activity leads to enhancer switching. **(A-D)** Overlap of the normalized 4C-seq signal between two conditions in a 300kb region surrounding the *Fos* viewpoint is shown. The y-axis is the number of relative reads normalized to 1M *cis* reads. The x-axis is the position along chr12 in Mb and extends from 150kb upstream to 150kb downstream of the *Fos* promoter viewpoint. (A) Shows the overlap (grey) between normalized 4C-seq signal in CON (red) and ETP rec. (green) neurons. The black boxes represent the regions that showed increased contact during ETP treatment. (B) Shows the overlap (grey) between normalized 4C-seq signal in CON (red) and ETP-NMDA (green) neurons. Arrow heads highlight *Fos’s* activity dependent enhancers E2 and E5 that incur changes in contact frequency. (C) Shows the overlap (grey) between normalized 4C-seq signal in ETP rec. (red) and ETP-NMDA (green) neurons. Arrow heads highlight *Fos’s* activity dependent enhancers E2 and E5 that incur changes in contact frequency. (D) Shows the overlap (grey) between normalized 4C-seq signal in NMDA (red) and ETP-NMDA (green) neurons. Arrow heads highlight *Fos’s* activity dependent enhancers E2 and E5 that incur changes in contact frequency. To annotate the chromatin regions/states surrounding the *Fos* promoter, we utilized the same reference tracks as in Figure S1. **(E-H)** Overlap of the normalized 4C-seq signal between two conditions in a 300kb region surrounding the *Npas4* viewpoint is shown. The y-axis is the number of relative reads normalized to 1M *cis* reads. The x-axis is the position along chr19 in Mb and extends from 150kb upstream to 150kb downstream of the *Npas4* promoter viewpoint. (F) Shows the overlap (grey) between normalized 4C-seq signal in CON (red) and ETP rec. (green) neurons. The black boxes represent the regions that showed increased contact during ETP treatment. (G) Shows the overlap (grey) between normalized 4C-seq signal in CON (red) and ETP-NMDA (green) neurons. (H) Shows the overlap (grey) between normalized 4C-seq signal in ETP rec. (red) and ETP-NMDA (green) neurons. Arrow head highlights an enhancer that only incurs contact increases in ETP-NMDA. (I) Shows the overlap (grey) between normalized 4C-seq signal in NMDA (red) and ETP-NMDA (green) neurons. Arrow head highlights an activity dependent enhancer that has contact increases in NMDA, but not in ETP-NMDA. To annotate the chromatin regions/states surrounding the *Npas4* promoter, we utilized previously published epigenetic datasets as shown in Figure S1.

**Figure S4:**
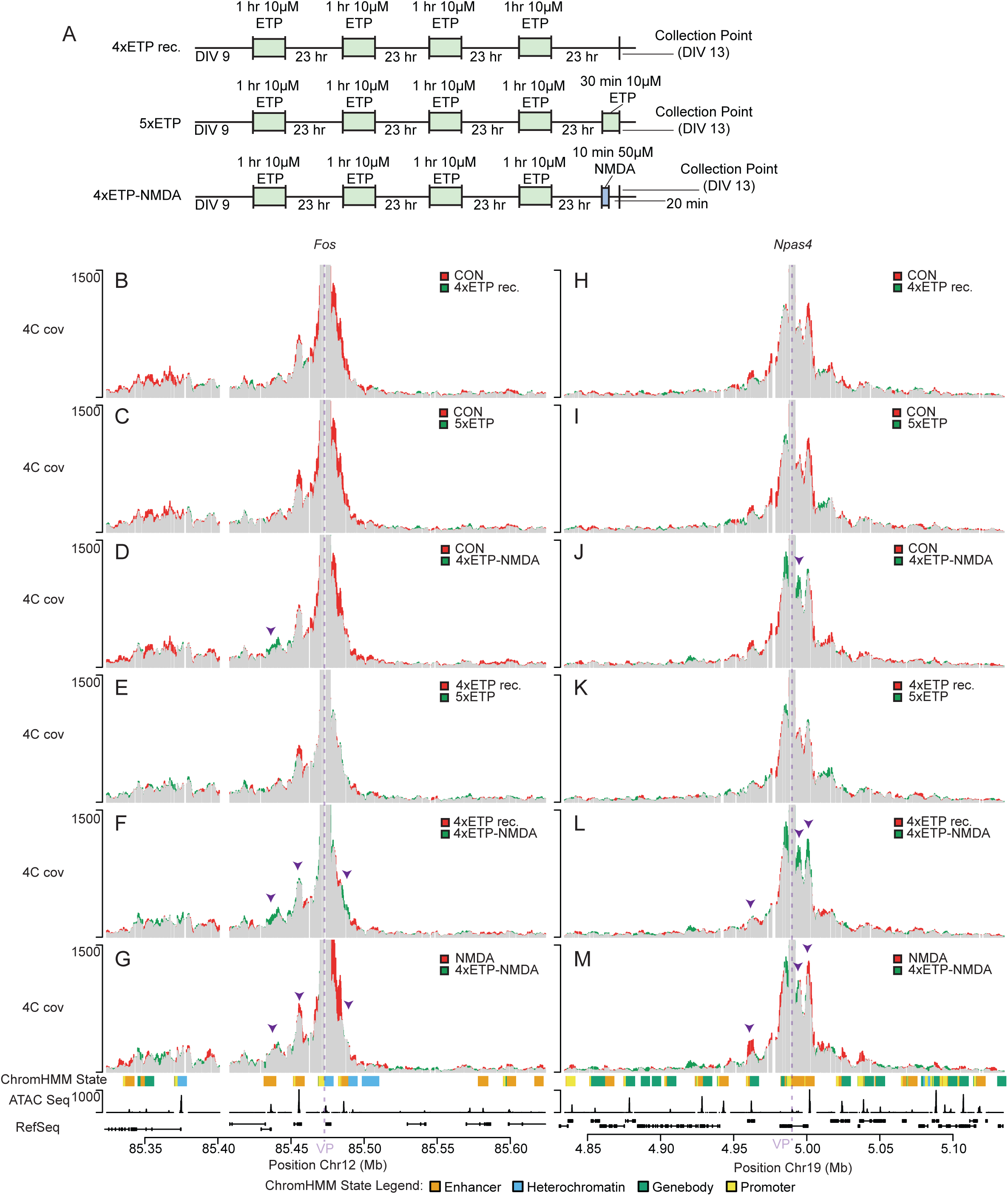
Multiple rounds of TOP2B-mediated break formation leads to novel enhancer contact increases and predominately decreases contacts along the cis chromosome. **(A)** Schematic diagram of the 4xETP rec., 5xETP, and 4xETP-NMDA treatments. DIV 9 neurons are stimulated with ETP (10μM, for 1 hr) followed by a ∼23 hr recovery period. This is followed by three more rounds of daily 1 hr ETP treatments followed by the recovery period. The neurons are then either collected after the final recovery period (4xETP rec.) on DIV 13 or after a fifth round of break formation is triggered by either a fifth ETP treatment (10M, 30min) (5xETP) or an NMDA treatment (50M, 10 min stimulation with a washout and a 20 min recovery period) (4xETP-NMDA). **(B-G)** Overlap of the normalized 4C-seq signal between two conditions in a 300kb region surrounding the *Fos* viewpoint is shown. The y-axis is the number of relative reads normalized to 1M *cis* reads. The x-axis is the position along chr12 in Mb and extends from 150kb upstream to 150kb downstream of the *Fos* promoter viewpoint. (B) Shows the overlap (grey) between normalized 4C-seq signal in CON (red) and 4xETP rec. (green) neurons. (C) Shows the overlap (grey) between normalized 4C-seq signal in CON (red) and 5xETP (green) neurons. (D) Shows the overlap (grey) between normalized 4C-seq signal in CON (red) and 4xETP-NMDA (green) neurons. Arrow head highlights *Fos’s* activity dependent enhancer E1 that only has gains in contact frequency after 4 rounds of previous TOP2B-mediated break formation. (E) Shows the overlap (grey) between normalized 4C-seq signal in 4xETP rec. (red) and 5xETP (green) neurons. (F) Shows the overlap (grey) between normalized 4C-seq signal in 4xETP rec. (red) and 4xETP-NMDA (green) neurons. Arrow heads highlight activity dependent *Fos* enhancers E1 (gained after 5 rounds of TOP2B-mediated break formation), E2 (contacted in ETP- and NMDA-treated neurons), and E5 (gained after 2 rounds of break formation). (G) Shows the overlap (grey) between normalized 4C-seq signal in NMDA (red) and 4xETP-NMDA (green) neurons. Arrow heads highlight activity dependent *Fos* enhancers E1 (gained after 5 rounds of TOP2B-mediated break formation), E2 (contacted in ETP- and NMDA-treated neurons), and E5 (gained after 2 rounds of break formation). Reference tracks are the same as those shown in Figure S1. **(H-M)** Overlap of the normalized 4C-seq signal between two conditions in a 300kb region surrounding the *Npas4* viewpoint is shown. The y-axis is the number of relative reads normalized to 1M *cis* reads. The x-axis is the position along chr19 in Mb and extends from 150kb upstream to 150kb downstream of the *Npas4* promoter viewpoint. (H) Shows the overlap (grey) between normalized 4C-seq signal in CON (red) and 4xETP rec. (green) neurons. (I) Shows the overlap (grey) between normalized 4C-seq signal in CON (red) and 5xETP (green) neurons. (J) Shows the overlap (grey) between normalized 4C-seq signal in CON (red) and 4xETP-NMDA (green) neurons. Arrow head highlights a putative internal enhancer that only has gains in contact frequency after 4 rounds of previous TOP2B-mediated break formation. (K) Shows the overlap (grey) between normalized 4C-seq signal in 4xETP rec. (red) and 5xETP (green) neurons. (L) Shows the overlap (grey) between normalized 4C-seq signal in 4xETP rec. (red) and 4xETP-NMDA (green) neurons. Arrow heads highlight activity dependent *Npas4* enhancers; the internal *Npas4* enhancer (gained after 5 rounds of TOP2B-mediated break formation), the enhancer downstream of *Npas4* TSS (contacted in ETP- and NMDA-treated neurons), and the enhancer upstream of *Npas4* TSS (gained after 2 rounds of break formation). (M) Shows the overlap (grey) between normalized 4C-seq signal in NMDA (red) and 4xETP-NMDA (green) neurons. Arrow heads highlight activity dependent *Npas4* enhancers; the internal *Npas4* enhancer (gained after 5 rounds of TOP2B-mediated break formation), the enhancer downstream of *Npas4* TSS (contacted in ETP- and NMDA-treated neurons), and the enhancer upstream of *Npas4* TSS (gained after 2 rounds of break formation). Reference tracks are the same as those shown in Figure S1.

**Figure S5:**
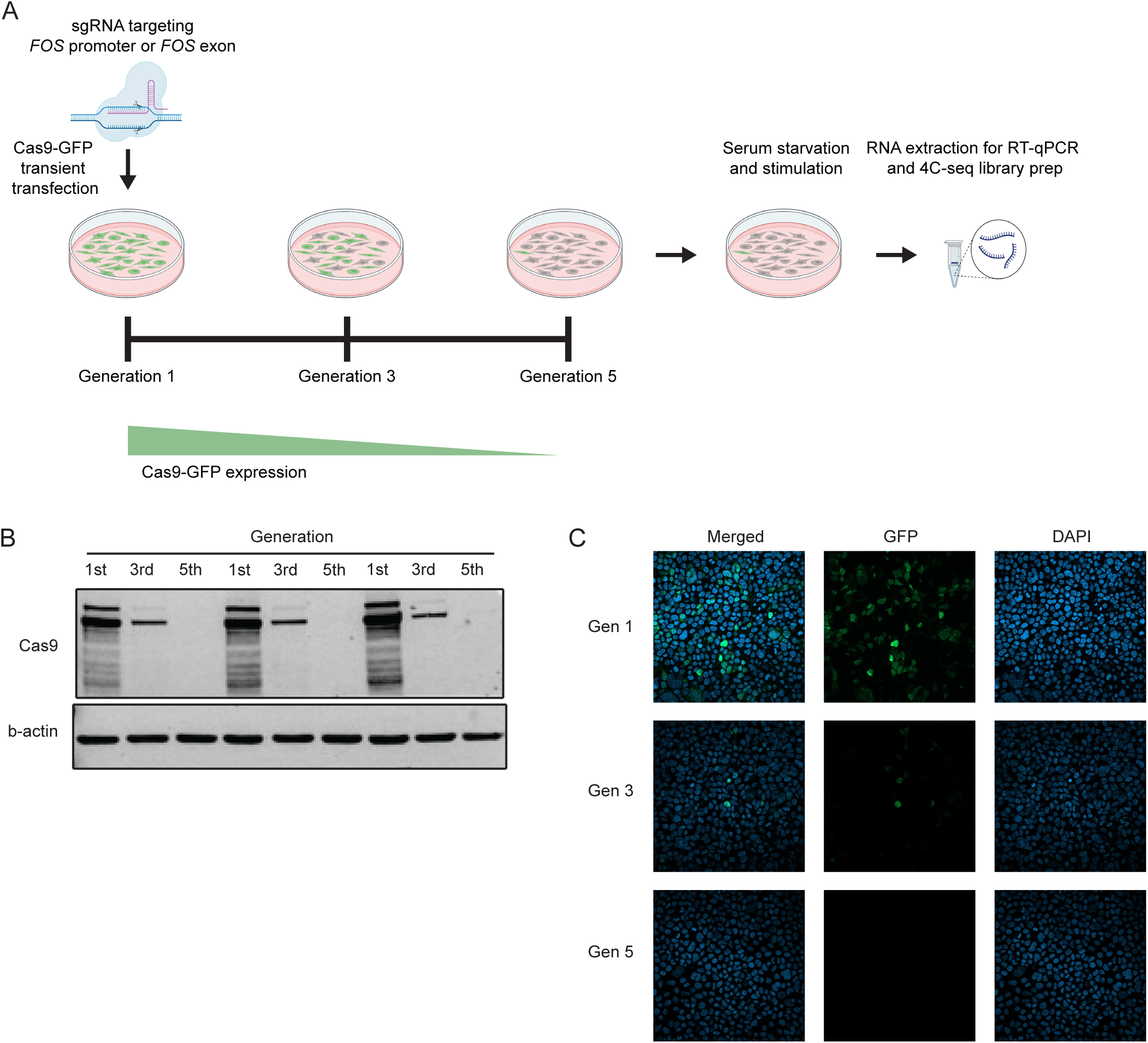
CRISPR Cas9 Recovery Experimental Paradigm and Validation in HEK293T cells. **(B)** Experimental schematic of CRISPR recovery and subsequent stimulation paradigm. HEK293T cells were transiently transfected with sgRNAs targeting either the promoter or exons of *FOS*. Generation 1 (Gen 1) cells were collected 96 hrs post transfection. Both CON and CRISPR HEK293T cells were then passaged for 5 generations until Cas9 expression was no longer detected (Gen 5). CON and CRISPR Gen 5 cells were then serum starved for ∼16.5 hrs and serum stimulated for 30 min prior to collection for either RNA extraction and RT-qPCR or 4C-seq library prep. **(C)** Western blot analysis of Cas9 and b-actin expression in CRISPR-Gen 1, CRISPR-Gen 3, and CRISPR-Gen 5 HEK293T cells **(D)** Confocal imaging of Cas9-GFP in CRISPR-Gen 1, CRISPR-Gen 3, and CRISPR-Gen 5 HEK293T cells.

## Notes

### Competing Interest Statement

The authors have declared no competing interest.

## REFERENCES

1. Flexner, J.B., Flexner, L.B., and Stellar, E. (1963). Memory in mice as affected by intracerebral puromycin. Science 141, 57–59. 10.1126/science.141.3575.57.

2. Goelet, P., Castellucci, V.F., Schacher, S., and Kandel, E.R. (1986). The long and the short of long-term memory--a molecular framework. Nature 322, 419–422. 10.1038/322419a0.

3. Nguyen, P.V., Abel, T., and Kandel, E.R. (1994). Requirement of a critical period of transcription for induction of a late phase of LTP. Science 265, 1104–1107. 10.1126/science.8066450.

4. Frey, U., Frey, S., Schollmeier, F., and Krug, M. (1996). Influence of actinomycin D, a RNA synthesis inhibitor, on long-term potentiation in rat hippocampal neurons in vivo and in vitro. J Physiol 490 (Pt 3), 703–711. 10.1113/jphysiol.1996.sp021179.

5. Yap, E.L., and Greenberg, M.E. (2018). Activity-Regulated Transcription: Bridging the Gap between Neural Activity and Behavior. Neuron 100, 330–348. 10.1016/j.neuron.2018.10.013.

6. West, A.E., and Greenberg, M.E. (2011). Neuronal activity-regulated gene transcription in synapse development and cognitive function. Cold Spring Harb Perspect Biol 3. 10.1101/cshperspect.a005744.

7. Madabhushi, R., and Kim, T.K. (2018). Emerging themes in neuronal activity-dependent gene expression. Mol Cell Neurosci 87, 27–34. 10.1016/j.mcn.2017.11.009.

8. Ebert, D.H., and Greenberg, M.E. (2013). Activity-dependent neuronal signalling and autism spectrum disorder. Nature 493, 327–337. 10.1038/nature11860.

9. Kim, T.K., Hemberg, M., Gray, J.M., Costa, A.M., Bear, D.M., Wu, J., Harmin, D.A., Laptewicz, M., Barbara-Haley, K., Kuersten, S., et al. (2010). Widespread transcription at neuronal activity-regulated enhancers. Nature 465, 182–187. 10.1038/nature09033.

10. Schaukowitch, K., Joo, J.Y., Liu, X., Watts, J.K., Martinez, C., and Kim, T.K. (2014). Enhancer RNA facilitates NELF release from immediate early genes. Mol Cell 56, 29–42. 10.1016/j.molcel.2014.08.023.

11. Fernandez-Albert, J., Lipinski, M., Lopez-Cascales, M.T., Rowley, M.J., Martin-Gonzalez, A.M., Del Blanco, B., Corces, V.G., and Barco, A. (2019). Immediate and deferred epigenomic signatures of in vivo neuronal activation in mouse hippocampus. Nat Neurosci 22, 1718–1730. 10.1038/s41593-019-0476-2.

12. Beagan, J.A., Pastuzyn, E.D., Fernandez, L.R., Guo, M.H., Feng, K., Titus, K.R., Chandrashekar, H., Shepherd, J.D., and Phillips-Cremins, J.E. (2020). Three-dimensional genome restructuring across timescales of activity-induced neuronal gene expression. Nat Neurosci 23, 707–717. 10.1038/s41593-020-0634-6.

13. Joo, J.Y., Schaukowitch, K., Farbiak, L., Kilaru, G., and Kim, T.K. (2016). Stimulus-specific combinatorial functionality of neuronal c-fos enhancers. Nat Neurosci 19, 75–83. 10.1038/nn.4170.

14. Sanborn, A.L., Rao, S.S., Huang, S.C., Durand, N.C., Huntley, M.H., Jewett, A.I., Bochkov, I.D., Chinnappan, D., Cutkosky, A., Li, J., et al. (2015). Chromatin extrusion explains key features of loop and domain formation in wild-type and engineered genomes. Proc Natl Acad Sci U S A 112, E6456–6465. 10.1073/pnas.1518552112.

15. Fudenberg, G., Imakaev, M., Lu, C., Goloborodko, A., Abdennur, N., and Mirny, L.A. (2016). Formation of Chromosomal Domains by Loop Extrusion. Cell Rep 15, 2038–2049. 10.1016/j.celrep.2016.04.085.

16. Rao, S.S.P., Huang, S.C., Glenn St Hilaire, B., Engreitz, J.M., Perez, E.M., Kieffer-Kwon, K.R., Sanborn, A.L., Johnstone, S.E., Bascom, G.D., Bochkov, I.D., et al. (2017). Cohesin Loss Eliminates All Loop Domains. Cell 171, 305–320 e324. 10.1016/j.cell.2017.09.026.

17. Davidson, I.F., Bauer, B., Goetz, D., Tang, W., Wutz, G., and Peters, J.M. (2019). DNA loop extrusion by human cohesin. Science 366, 1338–1345. 10.1126/science.aaz3418.

18. Kim, Y., Shi, Z., Zhang, H., Finkelstein, I.J., and Yu, H. (2019). Human cohesin compacts DNA by loop extrusion. Science 366, 1345–1349. 10.1126/science.aaz4475.

19. Bauer, B.W., Davidson, I.F., Canena, D., Wutz, G., Tang, W., Litos, G., Horn, S., Hinterdorfer, P., and Peters, J.M. (2021). Cohesin mediates DNA loop extrusion by a “swing and clamp” mechanism. Cell 184, 5448–5464 e5422. 10.1016/j.cell.2021.09.016.

20. Davidson, I.F., and Peters, J.M. (2021). Genome folding through loop extrusion by SMC complexes. Nat Rev Mol Cell Biol 22, 445–464. 10.1038/s41580-021-00349-7.

21. Calderon, L., Weiss, F.D., Beagan, J.A., Oliveira, M.S., Georgieva, R., Wang, Y.F., Carroll, T.S., Dharmalingam, G., Gong, W., Tossell, K., et al. (2022). Cohesin-dependence of neuronal gene expression relates to chromatin loop length. Elife 11. 10.7554/eLife.76539.

22. Madabhushi, R., Gao, F., Pfenning, A.R., Pan, L., Yamakawa, S., Seo, J., Rueda, R., Phan, T.X., Yamakawa, H., Pao, P.C., et al. (2015). Activity-Induced DNA Breaks Govern the Expression of Neuronal Early-Response Genes. Cell 161, 1592–1605. 10.1016/j.cell.2015.05.032.

23. Delint-Ramirez, I., Konada, L., Heady, L., Rueda, R., Jacome, A.S.V., Marlin, E., Marchioni, C., Segev, A., Kritskiy, O., Yamakawa, S., et al. (2022). Calcineurin dephosphorylates topoisomerase IIbeta and regulates the formation of neuronal-activity-induced DNA breaks. Mol Cell 82, 3794–3809 e3798. 10.1016/j.molcel.2022.09.012.

24. Bunch, H., Lawney, B.P., Lin, Y.F., Asaithamby, A., Murshid, A., Wang, Y.E., Chen, B.P., and Calderwood, S.K. (2015). Transcriptional elongation requires DNA break-induced signalling. Nat Commun 6, 10191. 10.1038/ncomms10191.

25. Pollina, E.A., Gilliam, D.T., Landau, A.T., Lin, C., Pajarillo, N., Davis, C.P., Harmin, D.A., Yap, E.L., Vogel, I.R., Griffith, E.C., et al. (2023). A NPAS4-NuA4 complex couples synaptic activity to DNA repair. Nature 614, 732–741. 10.1038/s41586-023-05711-7.

26. Stott, R.T., Kritsky, O., and Tsai, L.H. (2021). Profiling DNA break sites and transcriptional changes in response to contextual fear learning. PLoS One 16, e0249691. 10.1371/journal.pone.0249691.

27. Trotter, K.W., King, H.A., and Archer, T.K. (2015). Glucocorticoid Receptor Transcriptional Activation via the BRG1-Dependent Recruitment of TOP2beta and Ku70/86. Mol Cell Biol 35, 2799–2817. 10.1128/MCB.00230-15.

28. Wong, R.H., Chang, I., Hudak, C.S., Hyun, S., Kwan, H.Y., and Sul, H.S. (2009). A role of DNA-PK for the metabolic gene regulation in response to insulin. Cell 136, 1056–1072. 10.1016/j.cell.2008.12.040.

29. Ju, B.G., Lunyak, V.V., Perissi, V., Garcia-Bassets, I., Rose, D.W., Glass, C.K., and Rosenfeld, M.G. (2006). A topoisomerase IIbeta-mediated dsDNA break required for regulated transcription. Science 312, 1798–1802. 10.1126/science.1127196.

30. Suberbielle, E., Sanchez, P.E., Kravitz, A.V., Wang, X., Ho, K., Eilertson, K., Devidze, N., Kreitzer, A.C., and Mucke, L. (2013). Physiologic brain activity causes DNA double-strand breaks in neurons, with exacerbation by amyloid-beta. Nat Neurosci 16, 613–621. 10.1038/nn.3356.

31. Crewe, M., Segev, A., Rueda, R., and Madabhushi, R. (2024). Atypical Modes of CTCF Binding Facilitate Tissue-Specific and Neuronal Activity-Dependent Gene Expression States. Mol Neurobiol 61, 3240–3257. 10.1007/s12035-023-03762-5.

32. Ong, C.T., and Corces, V.G. (2014). CTCF: an architectural protein bridging genome topology and function. Nat Rev Genet 15, 234–246. 10.1038/nrg3663.

33. Simonis, M., Klous, P., Splinter, E., Moshkin, Y., Willemsen, R., de Wit, E., van Steensel, B., and de Laat, W. (2006). Nuclear organization of active and inactive chromatin domains uncovered by chromosome conformation capture-on-chip (4C). Nat Genet 38, 1348–1354. 10.1038/ng1896.

34. Krijger, P.H.L., Geeven, G., Bianchi, V., Hilvering, C.R.E., and de Laat, W. (2020). 4C-seq from beginning to end: A detailed protocol for sample preparation and data analysis. Methods 170, 17–32. 10.1016/j.ymeth.2019.07.014.

35. Delint-Ramirez, I., and Madabhushi, R. (2023). NPAS4 juggles neuronal activity-dependent transcription and DSB repair with NuA4. Mol Cell 83, 1208–1209. 10.1016/j.molcel.2023.03.019.

36. Delint-Ramirez, I., and Madabhushi, R. (2025). DNA damage and its links to neuronal aging and degeneration. Neuron 113, 7–28. 10.1016/j.neuron.2024.12.001.

37. Ernst, J., and Kellis, M. (2017). Chromatin-state discovery and genome annotation with ChromHMM. Nat Protoc 12, 2478–2492. 10.1038/nprot.2017.124.

38. Consortium, E.P., Moore, J.E., Purcaro, M.J., Pratt, H.E., Epstein, C.B., Shoresh, N., Adrian, J., Kawli, T., Davis, C.A., Dobin, A., et al. (2020). Expanded encyclopaedias of DNA elements in the human and mouse genomes. Nature 583, 699–710. 10.1038/s41586-020-2493-4.

39. Gorkin, D.U., Barozzi, I., Zhao, Y., Zhang, Y., Huang, H., Lee, A.Y., Li, B., Chiou, J., Wildberg, A., Ding, B., et al. (2020). An atlas of dynamic chromatin landscapes in mouse fetal development. Nature 583, 744–751. 10.1038/s41586-020-2093-3.

40. Segev, A., Heady, L., Crewe, M., and Madabhushi, R. (2024). Mapping catalytically engaged TOP2B in neurons reveals the principles of topoisomerase action within the genome. Cell Rep 43, 113809. 10.1016/j.celrep.2024.113809.

41. Raviram, R., Rocha, P.P., Muller, C.L., Miraldi, E.R., Badri, S., Fu, Y., Swanzey, E., Proudhon, C., Snetkova, V., Bonneau, R., and Skok, J.A. (2016). 4C-ker: A Method to Reproducibly Identify Genome-Wide Interactions Captured by 4C-Seq Experiments. PLoS Comput Biol 12, e1004780. 10.1371/journal.pcbi.1004780.

42. Dixon, J.R., Selvaraj, S., Yue, F., Kim, A., Li, Y., Shen, Y., Hu, M., Liu, J.S., and Ren, B. (2012). Topological domains in mammalian genomes identified by analysis of chromatin interactions. Nature 485, 376–380. 10.1038/nature11082.

43. Mirny, L.A., Imakaev, M., and Abdennur, N. (2019). Two major mechanisms of chromosome organization. Curr Opin Cell Biol 58, 142–152. 10.1016/j.ceb.2019.05.001.

44. Lieberman-Aiden, E., van Berkum, N.L., Williams, L., Imakaev, M., Ragoczy, T., Telling, A., Amit, I., Lajoie, B.R., Sabo, P.J., Dorschner, M.O., et al. (2009). Comprehensive mapping of long-range interactions reveals folding principles of the human genome. Science 326, 289–293. 10.1126/science.1181369.

45. Bonev, B., Mendelson Cohen, N., Szabo, Q., Fritsch, L., Papadopoulos, G.L., Lubling, Y., Xu, X., Lv, X., Hugnot, J.P., Tanay, A., and Cavalli, G. (2017). Multiscale 3D Genome Rewiring during Mouse Neural Development. Cell 171, 557–572 e524. 10.1016/j.cell.2017.09.043.

46. Durand, N.C., Shamim, M.S., Machol, I., Rao, S.S., Huntley, M.H., Lander, E.S., and Aiden, E.L. (2016). Juicer Provides a One-Click System for Analyzing Loop-Resolution Hi-C Experiments. Cell Syst 3, 95–98. 10.1016/j.cels.2016.07.002.

47. Brinkman, E.K., Chen, T., de Haas, M., Holland, H.A., Akhtar, W., and van Steensel, B. (2018). Kinetics and Fidelity of the Repair of Cas9-Induced Double-Strand DNA Breaks. Mol Cell 70, 801–813 e806. 10.1016/j.molcel.2018.04.016.

48. Allen, F., Crepaldi, L., Alsinet, C., Strong, A.J., Kleshchevnikov, V., De Angeli, P., Palenikova, P., Khodak, A., Kiselev, V., Kosicki, M., et al. (2018). Predicting the mutations generated by repair of Cas9-induced double-strand breaks. Nat Biotechnol. 10.1038/nbt.4317.

49. Harmsen, T.J.W., Pritchard, C.E.J., Riepsaame, J., van de Vrugt, H.J., Huijbers, I.J., and Te Riele, H. (2022). HideRNAs protect against CRISPR-Cas9 re-cutting after successful single base-pair gene editing. Sci Rep 12, 9606. 10.1038/s41598-022-13688-y.

50. Rajagopal, N., Srinivasan, S., Kooshesh, K., Guo, Y., Edwards, M.D., Banerjee, B., Syed, T., Emons, B.J., Gifford, D.K., and Sherwood, R.I. (2016). High-throughput mapping of regulatory DNA. Nat Biotechnol 34, 167–174. 10.1038/nbt.3468.

51. Pradeepa, M.M., Grimes, G.R., Kumar, Y., Olley, G., Taylor, G.C., Schneider, R., and Bickmore, W.A. (2016). Histone H3 globular domain acetylation identifies a new class of enhancers. Nat Genet 48, 681–686. 10.1038/ng.3550.

52. Schoenfelder, S., and Fraser, P. (2019). Long-range enhancer-promoter contacts in gene expression control. Nat Rev Genet 20, 437–455. 10.1038/s41576-019-0128-0.

53. Magnitov, M., and de Wit, E. (2024). Attraction and disruption: how loop extrusion and compartmentalisation shape the nuclear genome. Curr Opin Genet Dev 86, 102194. 10.1016/j.gde.2024.102194.

54. Gu, B., Swigut, T., Spencley, A., Bauer, M.R., Chung, M., Meyer, T., and Wysocka, J. (2018). Transcription-coupled changes in nuclear mobility of mammalian cis-regulatory elements. Science 359, 1050–1055. 10.1126/science.aao3136.

55. Dimitrova, N., Chen, Y.C., Spector, D.L., and de Lange, T. (2008). 53BP1 promotes non-homologous end joining of telomeres by increasing chromatin mobility. Nature 456, 524–528. 10.1038/nature07433.

56. Krawczyk, P.M., Borovski, T., Stap, J., Cijsouw, T., ten Cate, R., Medema, J.P., Kanaar, R., Franken, N.A., and Aten, J.A. (2012). Chromatin mobility is increased at sites of DNA double-strand breaks. J Cell Sci 125, 2127–2133. 10.1242/jcs.089847.

57. Marnef, A., and Legube, G. (2017). Organizing DNA repair in the nucleus: DSBs hit the road. Curr Opin Cell Biol 46, 1–8. 10.1016/j.ceb.2016.12.003.

58. Seeber, A., Hauer, M.H., and Gasser, S.M. (2018). Chromosome Dynamics in Response to DNA Damage. Annu Rev Genet 52, 295–319. 10.1146/annurev-genet-120417-031334.

59. Tehrani, S.S.H., Kogan, A., Mikulski, P., and Jansen, L.E.T. (2025). Remembering foods and foes: emerging principles of transcriptional memory. Cell Death Differ 32, 16–26. 10.1038/s41418-023-01200-6.

60. Su, Y., Shin, J., Zhong, C., Wang, S., Roychowdhury, P., Lim, J., Kim, D., Ming, G.L., and Song, H. (2017). Neuronal activity modifies the chromatin accessibility landscape in the adult brain. Nat Neurosci 20, 476–483. 10.1038/nn.4494.

61. Marco, A., Meharena, H.S., Dileep, V., Raju, R.M., Davila-Velderrain, J., Zhang, A.L., Adaikkan, C., Young, J.Z., Gao, F., Kellis, M., and Tsai, L.H. (2020). Mapping the epigenomic and transcriptomic interplay during memory formation and recall in the hippocampal engram ensemble. Nat Neurosci 23, 1606–1617. 10.1038/s41593-020-00717-0.

62. Engmann, O., Labonte, B., Mitchell, A., Bashtrykov, P., Calipari, E.S., Rosenbluh, C., Loh, Y.E., Walker, D.M., Burek, D., Hamilton, P.J., et al. (2017). Cocaine-Induced Chromatin Modifications Associate With Increased Expression and Three-Dimensional Looping of Auts2. Biol Psychiatry 82, 794–805. 10.1016/j.biopsych.2017.04.013.

63. Zhao, Z., Zhang, Z., Li, J., Dong, Q., Xiong, J., Li, Y., Lan, M., Li, G., and Zhu, B. (2020). Sustained TNF-alpha stimulation leads to transcriptional memory that greatly enhances signal sensitivity and robustness. Elife 9. 10.7554/eLife.61965.

64. Iberg-Badeaux, A., Collombet, S., Laurent, B., van Oevelen, C., Chin, K.K., Thieffry, D., Graf, T., and Shi, Y. (2017). A Transcription Factor Pulse Can Prime Chromatin for Heritable Transcriptional Memory. Mol Cell Biol 37. 10.1128/MCB.00372-16.

65. Arnould, C., Rocher, V., Saur, F., Bader, A.S., Muzzopappa, F., Collins, S., Lesage, E., Le Bozec, B., Puget, N., Clouaire, T., et al. (2023). Chromatin compartmentalization regulates the response to DNA damage. Nature. 10.1038/s41586-023-06635-y.

66. Delint-Ramirez, I., Konada, L., Heady, L., Rueda, R., Jacome, A.S.V., Marlin, E., Marchioni, C., Segev, A., Kritskiy, O., Yamakawa, S., et al. (2022). Calcineurin dephosphorylates topoisomerase IIbeta and regulates the formation of neuronal-activity-induced DNA breaks. Mol Cell. 10.1016/j.molcel.2022.09.012.

67. Geeven, G., Teunissen, H., de Laat, W., and de Wit, E. (2018). peakC: a flexible, non-parametric peak calling package for 4C and Capture-C data. Nucleic Acids Res 46, e91. 10.1093/nar/gky443.

68. Quinlan, A.R. (2014). BEDTools: The Swiss-Army Tool for Genome Feature Analysis. Curr Protoc Bioinformatics 47, 11 12 11–34. 10.1002/0471250953.bi1112s47.

69. Quinlan, A.R., and Hall, I.M. (2010). BEDTools: a flexible suite of utilities for comparing genomic features. Bioinformatics 26, 841–842. 10.1093/bioinformatics/btq033.

70. Ernst, J., and Kellis, M. (2012). ChromHMM: automating chromatin-state discovery and characterization. Nat Methods 9, 215–216. 10.1038/nmeth.1906.

